# Tissue Context Shapes the Circadian Transcriptome of the Arabidopsis Leaf

**DOI:** 10.1101/2025.06.12.659411

**Authors:** Alveena Zulfiqar, Zachary A. Myers, Danielle Schoenecker, Ananda Menon, Kathleen Greenham

## Abstract

Circadian regulation enables plants to coordinate cellular processes with daily environmental cycles, yet how clocks operate across cell and tissue types remains poorly understood. To resolve circadian regulation in the mature *Arabidopsis thaliana* leaf, we generated a 24-hour single nucleus RNA-sequencing (snRNA-seq) circadian time course spanning seven circadian time points and ∼30,000 nuclei. We recovered all major leaf cell types and identified >7,400 genes with cluster-resolved rhythmic expression. Coexpression analyses defined five major temporal expression clades, revealing cell-type-specific phase shifts, while cluster-specific gene regulatory networks identified shared and unique transcription factor targets. These analyses suggest that core clock components broadly coordinate processes such as photosynthesis and light signaling while fine-tuning hormone, redox, and stress pathways in cell-type-specific manners. To restore the spatial information lost during nuclei isolation, we designed a 100-gene Xenium spatial transcriptomics panel and profiled intact mature leaves at relative dawn and dusk. Spatial transcriptomics validated cell-type-associated expression patterns from the single-nucleus time course and further revealed spatially restricted domains that were not apparent from cluster averages alone, including vascular subdomains and guard cell-associated differences in clock transcript accumulation. Several core clock components had unequal spatial distribution across similar cell types, suggesting potential fine-tuning of the clock in discrete spatial regions. Together, these findings show that the circadian transcriptional programs in the mature leaf are not simply cell-type-specific but spatially organized within the architecture of the organ.

**Significance:** Plants lack an animal-like central circadian pacemaker, raising the question of how local clocks are organized and coordinated across tissues. Here, we asked how circadian timing and transcriptional control vary among cell types by combining a 24-hour single-nucleus RNA-sequencing time course, cell type-resolved gene regulatory network inference, and Xenium spatial transcriptomics in mature Arabidopsis leaves. We identified widespread rhythmic expression across leaf cell types and uncovered shared and cell type-specific regulatory connections for core clock transcription factors. Spatial profiling preserved tissue organization and revealed heterogeneous clock transcript accumulation that was not fully resolved by dissociation-based measurements. These findings establish tissue context as a major determinant of how the plant clock coordinates metabolism, growth, and environmental responses.

## Introduction

The Earth’s rotation on its axis generates predictable fluctuations in the day-night cycle. Over time, organisms from unicellular to complex multicellular species, including humans, have evolved endogenous circadian oscillators (Saini et al. 2019). These autonomous time-keeping mechanisms allow organisms to anticipate and adapt to daily changes in light and temperature (Saini et al. 2019; Micklem and Locke 2021). The circadian mediated alignment of cellular processes with external cues is central to survival and fitness (Ouyang et al. 1998). Although clock genes vary across kingdoms, the fundamental principles of circadian regulation—such as transcriptional-translational feedback loops and entrainment by environmental cues known as zeitgebers—are remarkably conserved (Achermann and Kunz 1999; Glossop and Hardin 2002; Hastings et al. 2003; Akashi and Takumi 2005; Adams et al. 2015).

In plants, the circadian clock is integral to coordinating physiological processes with daily environmental changes, ensuring optimal growth, metabolism and development (Dodd et al. 2005; Graf et al. 2010; Greenham and McClung 2015). The plant circadian system consists of transcriptional regulators that function through transcriptional-translational feedback loops to generate rhythmic gene expression patterns within a roughly 24-hour period (Pokhilko et al. 2012; Nohales and Kay 2016). Dawn expressed *CIRCADIAN CLOCK ASSOCIATED 1* (*CCA1*) and *LATE ELONGATED HYPOCOTYL* (*LHY*) MYB transcription factors (TFs) are followed by the sequential expression of the transcriptional repressors *PSEUDO RESPONSE REGULATORS PRR9, PRR7* and *PRR5* and the evening-phased *TIMING OF CAB EXPRESSION 1 (TOC1)/PRR1 (Schaffer et al. 1998; Wang and Tobin 1998; Strayer et al. 2000; Alabadí et al. 2001; Nakamichi et al. 2010; Xie et al. 2014)*. Additional repressors include the Evening Complex (EC), consisting of LUX ARRHYTHMO (LUX)-EARLY FLOWERING 3 (ELF3) and ELF4 that contribute to circadian clock function and coordinating growth and development (Nusinow et al. 2011). NIGHT LIGHT-INDUCIBLE AND CLOCK-REGULATED1 (LNK1) and LNK2 act as co-activators of REVEILLE4 (RVE4) and RVE8 to positively regulate *PRR5, TOC1* and the EC (Farinas and Mas 2011; Rawat et al. 2011; Rugnone et al. 2013; Xie et al. 2014). These regulatory arrangements facilitate robust circadian gene expression across environmental variation and allow the clock to gate and influence target genes across time and tissue types.

Unlike mammals, whose suprachiasmatic nucleus is the master pacemaker (Ma and Elizabeth 2019), plants have decentralized tissue and organ oscillators (Gould et al. 2018), with evidence of hierarchical organization. The shoot apical meristem (SAM) can act as a dominant clock (Takahashi et al. 2015) and signal to roots (Ikeda et al. 2023), yet detached leaves and roots remain rhythmic (Li et al. 2020), showing that sustained oscillation does not require the SAM. Local clocks also control distinct processes: the vascular clock regulates flowering through FLOWERING LOCUS T (FT) in phloem companion cells (Endo et al. 2014; Shimizu et al. 2015; Voß et al. 2015), epidermal clocks regulate temperature-dependent elongation (Shimizu et al. 2015), and vascular signals can dominate mesophyll rhythms in leaves (Endo et al. 2014; Shimizu et al. 2015; Voß et al. 2015).

Plant organs and cell types maintain distinct circadian periods (Thain et al. 2000; Fukuda et al. 2007; James et al. 2008; Wenden et al. 2012; Endo et al. 2014; Gould et al. 2018; Greenwood et al. 2019). In Arabidopsis seedlings, cotyledons and hypocotyls cycle faster than roots (James et al. 2008; Gould et al. 2018; Greenwood et al. 2019), guard cells more slowly than adjacent mesophyll and epidermal cells (Yakir et al. 2011), and vascular tissues more robustly than mesophyll (James et al. 2008; Endo et al. 2014). Coupling also varies spatially: dense shoot and root meristems are more synchronized, whereas leaves sustain independent phases of clock-gene promoter activity (Fukuda et al. 2007; Wenden et al. 2012; Takahashi et al. 2015). Near-cellular pCCA1::CCA1-YFP and GIGANTEA promoter-driven LUCIFERASE (GI::LUC) imaging similarly showed shorter periods in root tips than elsewhere in the root (Gould et al. 2018; Greenwood et al. 2019). These findings establish intra-tissue clock partitioning, but promoter reporters and bulk sampling cannot resolve the spatial domains of clock and clock-regulated transcripts, requiring methods that preserve cellular identity and tissue architecture.

Spatial transcriptomics addresses this gap by measuring transcripts in intact tissue. Plant methods include spatial barcoding platforms such as Visium (Ståhl et al. 2016) and Stereo-seq (Chen et al. 2022), and imaging-based RNAscope (Wang et al. 2012), MERFISH (Chen et al. 2015), and Xenium (Janesick et al. 2023). Applications span development, stress, cell identity, and cross-species comparisons (Nobori 2025; Long et al. 2026; Peirats-Llobet et al. 2026), despite challenges from plant anatomy and autofluorescence. In circadian studies, these technologies can test whether single-cell or single-nucleus temporal patterns persist in situ and reveal heterogeneity, neighborhood relationships, and localized expression lost through dissociation.

Bulk RNA sequencing (RNA-seq) and microarrays obscure cell-specific rhythms by averaging heterogeneous tissues. Single-cell and single-nucleus RNA-sequencing now resolves cell-specific circadian regulation (Lopez-Anido et al. 2021; Shahan et al. 2022; Guo et al. 2025). An snRNA-seq time course of 8-day-old Arabidopsis seedlings found core clock genes rhythmic across most cell types but variable in phase and amplitude (Qin et al. 2025). Most rhythmic genes were cluster-specific; cotyledon subclusters shared more cycling genes than root subclusters, with phase differences even within cell types (Apelt et al. 2022; Qin et al. 2025). Integrating single-nucleus and spatial transcriptomics can define how these oscillators regulate mature-plant processes and determine whether circadian programs reflect cell identity alone or also position within the mature leaf, informing targeted growth improvement.

In this study, we captured the transcriptional profiles of over 30,000 nuclei across a 24 hour circadian time course to investigate circadian regulation in mature leaves of Arabidopsis, and leveraged these observations to generate targeted circadian spatial, 100-gene transcriptomic profiles of the mature Arabidopsis leaf. This approach revealed widespread rhythmic transcription across major leaf cell types, including robust cycling of core oscillator components, while also uncovering cell-type-specific differences in rhythmic gene expression, phase structure, and downstream clock-associated outputs. Temporal co-expression analysis identified major phase-dependent expression programs shared across the leaf, and cluster-resolved gene regulatory networks identified both conserved and cell-type-restricted regulatory modules associated with core clock regulators, including CCA1, LNK2, and PRR7. Xenium spatial profiling supported the cell type and time of day dependent expression of many of these targets, and further uncovered spatial heterogeneity in the accumulation of many clock TFs, including *CCA1, TOC1,* and *PRR7*. Together, these analyses show that the mature leaf contains a broadly distributed circadian system whose phase and transcriptional outputs are shaped by cell identity and spatial context, providing a framework for dissecting decentralized clock architecture and its role in coordinating tissue-level physiology.

## Results

### Construction and cell type identification of a circadian single nucleus time course in mature leaves of Arabidopsis

To resolve circadian regulation across Arabidopsis leaf cell types, we performed a single-nucleus RNA-sequencing (snRNA-seq) time course in 13-day-old plants. Seedlings were entrained under a 12-hour photoperiod at 22°C for 11 days, then transferred to constant light (LL) (Fig. 1A). Duplicate above-ground samples were collected every four hours for 24 hours beginning at ZT48 (zeitgeber time, hours after dawn) by cutting at the root:shoot junction, and fresh tissue was immediately processed for nuclei isolation and 10X Genomics capture. Cellranger assessment (Zheng et al. 2017) showed ∼2,000–4,000 nuclei per library and median detection of ∼1,000–1,500 genes, comparable to previous plant single-nucleus libraries (*Dataset S1*) (Wang et al. 2023; Guo et al. 2025; Qin et al. 2025). Background RNA was higher than expected, likely because we minimized handling by omitting fluorescence-activated nuclei sorting and density-gradient purification. Of 14 libraries, ZT24B failed QC and ZT04B had reduced nuclei recovery (*Dataset S1*).

**Figure 1.**
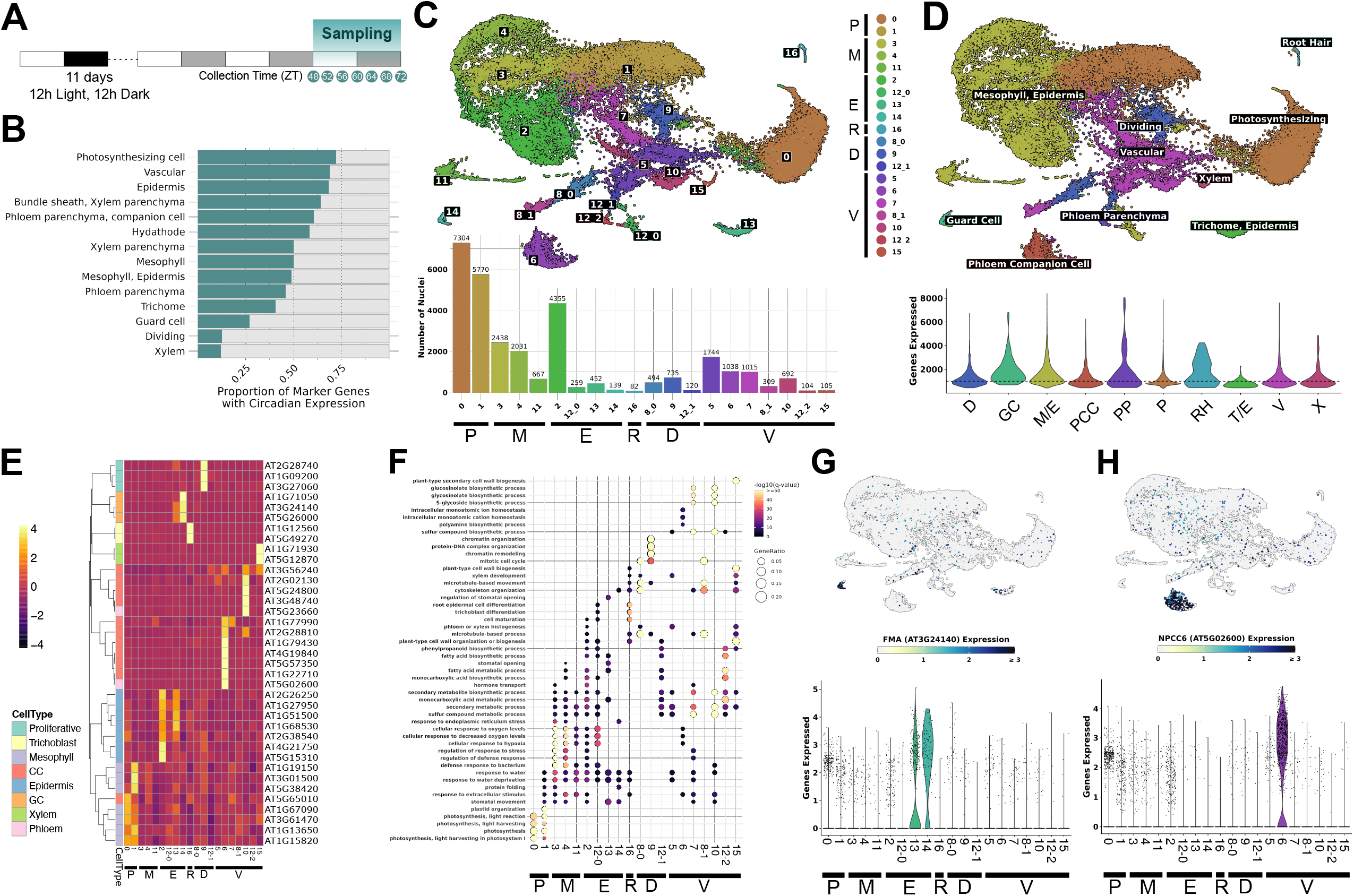
Leaf cell types across a circadian time course. (A) Growth and sampling design. (B) Circadian-regulated cell-type markers. (C) Cycling-masked integrated UMAP and nuclei per cluster. (D) Cell-type UMAP and genes detected per group. (E) Scaled marker expression. (F) Gene Ontology (GO) enrichments of cluster biomarkers. (G–H) *FMA* (AT3G24140; guard-cell c14) and *NPCC6* (AT5G02600; phloem companion-cell c6) feature plots. D, dividing; E, epidermis; GC, guard cell; M, mesophyll; M/E, mesophyll/epidermis; P, photosynthesizing; PCC, phloem companion cell; PP, phloem parenchyma; R, root; RH, root hair; T/E, trichome/epidermis; V, vascular; X, xylem.

Unguided integration of time-course data can produce phase-specific clusters when time-of-day genes obscure cell type-specific signatures (Wen et al. 2020; Qin et al. 2025). Because broadly expressed clock components such as CCA1 may introduce such biases (Gould et al. 2018), we examined marker genes from previous studies. Many cluster-specific markers were circadian regulated (Fig. 1B), and several cell types were defined predominantly by cycling genes (Romanowski et al. 2020; Lee et al. 2023). Masking ∼7,000 circadian genes during anchor selection and using 15,000 variable genes as integration anchors yielded cell type-specific clustering (Fig. 1C). The masked list combined cycling genes from bulk RNA-seq (Romanowski et al. 2020; Lee et al. 2023) with genes rhythmic in our pseudo-bulk data (*SI Appendix*). The resulting dataset contained ∼30,000 high-quality nuclei representing all major Arabidopsis leaf cell types. Most were photosynthetic or mesophyll nuclei, with fewer guard, phloem companion, and dividing cells (Fig. 1D). These included cell types rich in circadian markers, such as guard cells, that were often unresolved without masking.

Assignments were supported by published markers (*Dataset S2*) and refined with Gene Ontology (GO) enrichment and cluster biomarkers (Fig. 1E-H, *Dataset S3*). Marker expression identified Clusters 6, 9, and 14 as phloem companion, dividing, and guard cells, respectively (Fig. 1E). Enriched GO terms also distinguished mesophyll, epidermal, and photosynthetic origins through water-use, fatty-acid-biosynthesis, and light-harvesting functions (Fig. 1F, *Dataset S1*). Cluster biomarkers further refined annotations (Fig. S1). *FAMA*, a bHLH regulator of stomatal development (Bergmann et al. 2004), was highly specific to Cluster 14, with moderate expression only in Cluster 13 (Fig. 1G). *NPCC6* (*NUCLEAR-ENRICHED PHLOEM COMPANION CELL GENE 6*), previously reported as phloem-companion-cell specific (Zhang et al. 2008), was a significant Cluster 6 biomarker (Fig. 1H). Together, these analyses identified all major leaf cell types across the circadian snRNA-seq time course.

### Cycling gene expression is widespread across all cell types of the mature Arabidopsis leaf

To assess circadian regulation, we examined pseudo-bulk profiles of selected clock components and outputs across the single-nucleus time course (Fig. 2A). All showed robust oscillations consistent with previous reports (Harmer et al. 2000; Covington et al. 2008; Romanowski et al. 2020). Cluster-resolved analysis (Fig. 2B, S2) revealed broadly similar *CCA1* and *TOC1* rhythms across many clusters, whereas other components cycled only in subsets. *GI*, *LUX*, and *PRR5* generally followed consistent patterns wherever they cycled, but several components shifted phase in specific clusters, including an earlier *ELF3* phase in guard cells (Fig. 2B). We used JTKcycle (Hughes et al. 2010) to identify cycling genes from pseudo-bulk and cluster-resolved expression at adjusted *P* < 0.05. This identified 4,769 cycling genes in the pseudo-bulk dataset (*Dataset S4*), of which ∼68% (3,264/4,769) overlapped a previous bulk RNA-seq circadian gene list (Harmer et al. 2000; Covington et al. 2008; Romanowski et al. 2020).

**Figure 2.**
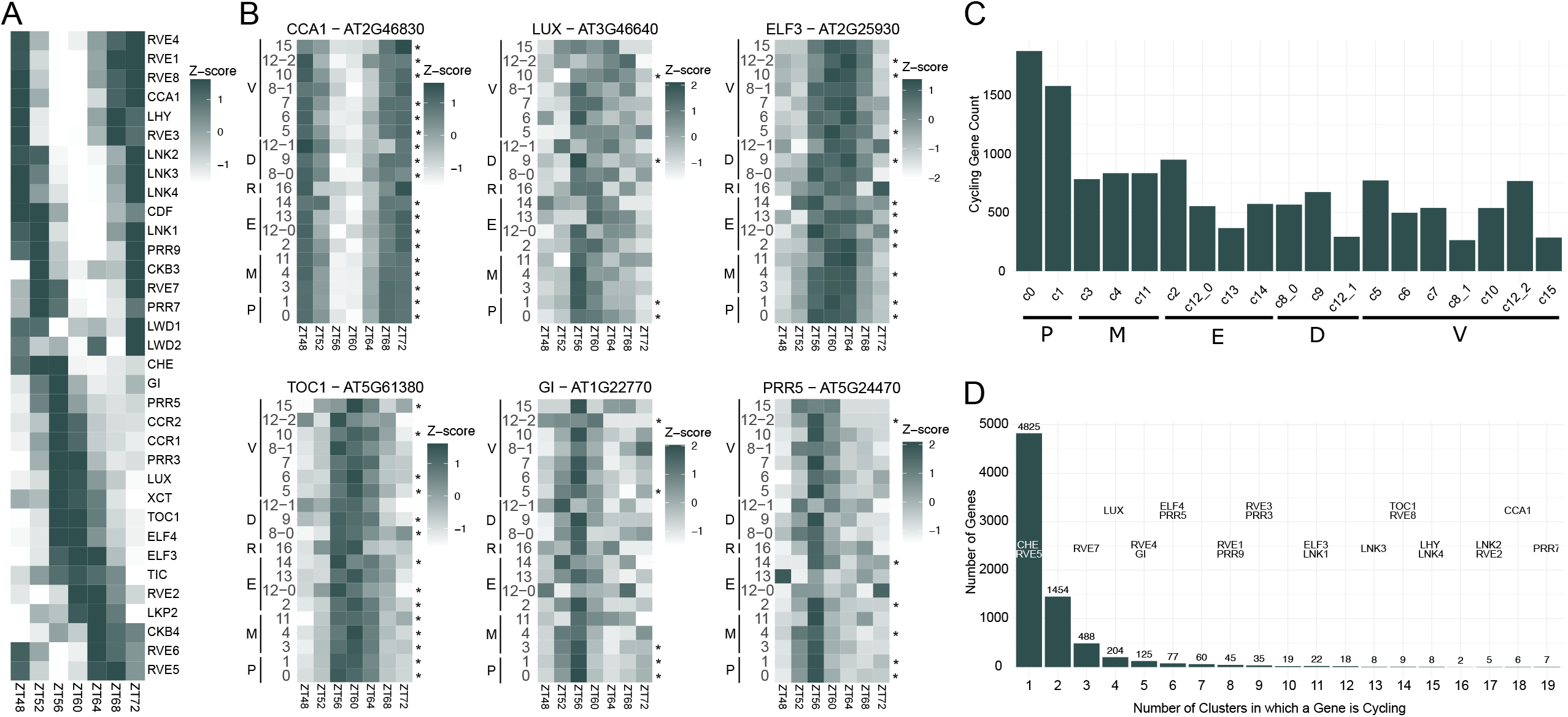
Circadian expression across leaf cell types. (A) Pseudobulk clock-gene and output expression. (B) Cluster-resolved *CCA1, TOC1, LUX, GI, ELF3,* and *PRR5* expression; asterisks mark significant cycling. (C) CycleJTK rhythmic genes per cluster. (D) Rhythmic genes grouped by the number of cycling clusters; key clock TFs are indicated.

Cluster-specific cycling genes were detected in every cluster except Cluster 16, where variable sampling likely caused uneven recovery of root hair cells across time points (Fig. S3). Counts ranged from ∼1,800 genes in Cluster 0 to ∼300 in Cluster 8_1 (Fig. 2D, *Dataset S5*). Although smaller clusters generally yielded fewer cycling genes, robust rhythms were consistently recovered for known clock TFs such as CCA1 and TOC1 (Fig. 2B). Overall, 4,825 genes cycled in at least one cluster, and more than half cycled in only one (Fig. 2E). Most clusters contained genes peaking throughout the time course, although some showed phase biases, including enrichment of morning-phased genes in guard cells (Fig. S4). *CCA1* and *TOC1* cycled significantly in 18 and 14 clusters, respectively, and the guard-cell *ELF3* phase shift was significant (Fig. 2B). Processing fresh nuclei at every time point precluded a 48-hour time course; therefore, we included biological replicates at each time point and applied a stringent cycling threshold to limit false negatives from sampling one cycle (Hughes et al. 2017).

Across clusters, we identified 7,417 rhythmic genes, 59% of which overlapped previous bulk RNA-seq data (Fig. S5) (Harmer et al. 2000; Covington et al. 2008; Romanowski et al. 2020). Comparison with a recent 24- and 48-hour circadian snRNA-seq dataset (Qin et al. 2025) showed 41% overlap with the 24-hour dataset and 63% with the 48-hour dataset (Fig. S5). The stronger 48-hour overlap supports our replicate design and indicates that our analysis recovered many genes previously identified as circadian regulated in bulk and single-cell studies.

### Coexpression network analysis identifies five major temporal expression patterns across all cell types

Due to the limitations inherent in calling phase from a single 24-hour cycle (Hughes et al. 2010), we confirmed gene cycling and phase using the weighted gene correlation network analysis (WGCNA) package in R (Langfelder and Horvath 2008). The method has been previously applied for grouping cycling genes with similar phases into modules (Greenham et al. 2017, 2020). This resulted in clustering of genes based on expression pattern into 26 modules. To remove genes with weak correlations with the module eigengene pattern, we filtered the modules by kME (Module Eigengene connectivity) value; only genes with kME ≥ 0.6 were retained. We next calculated correlation metrics for the modules identified by using the average zScore of each module for the time course. We then used the metrics to perform hierarchical clustering of the modules. Euclidean distance calculated from the correlation values, where shorter distance indicates strong relationship and more similarity between the modules, identified five major clades among all modules. Representative expression patterns for each clade was calculated by averaging the zScore of all the genes and resulted in five major gene expression patterns throughout the day: early morning genes (clade 3), morning genes (clade 2), mid-day genes (clade 1), evening genes (clade 4) and late-night genes (clade 5) (Fig. 3A, *Dataset S6*). Additionally, we examined the distribution of the five clades across our clusters, and each cell type had representation of all five expression patterns (Fig. 3B). Some cell types were more enriched in certain expression patterns, for example C5 (Vasculature) and C14 (GC) clusters had a larger proportion of genes with morning expression pattern (clade 2).

**Figure 3.**
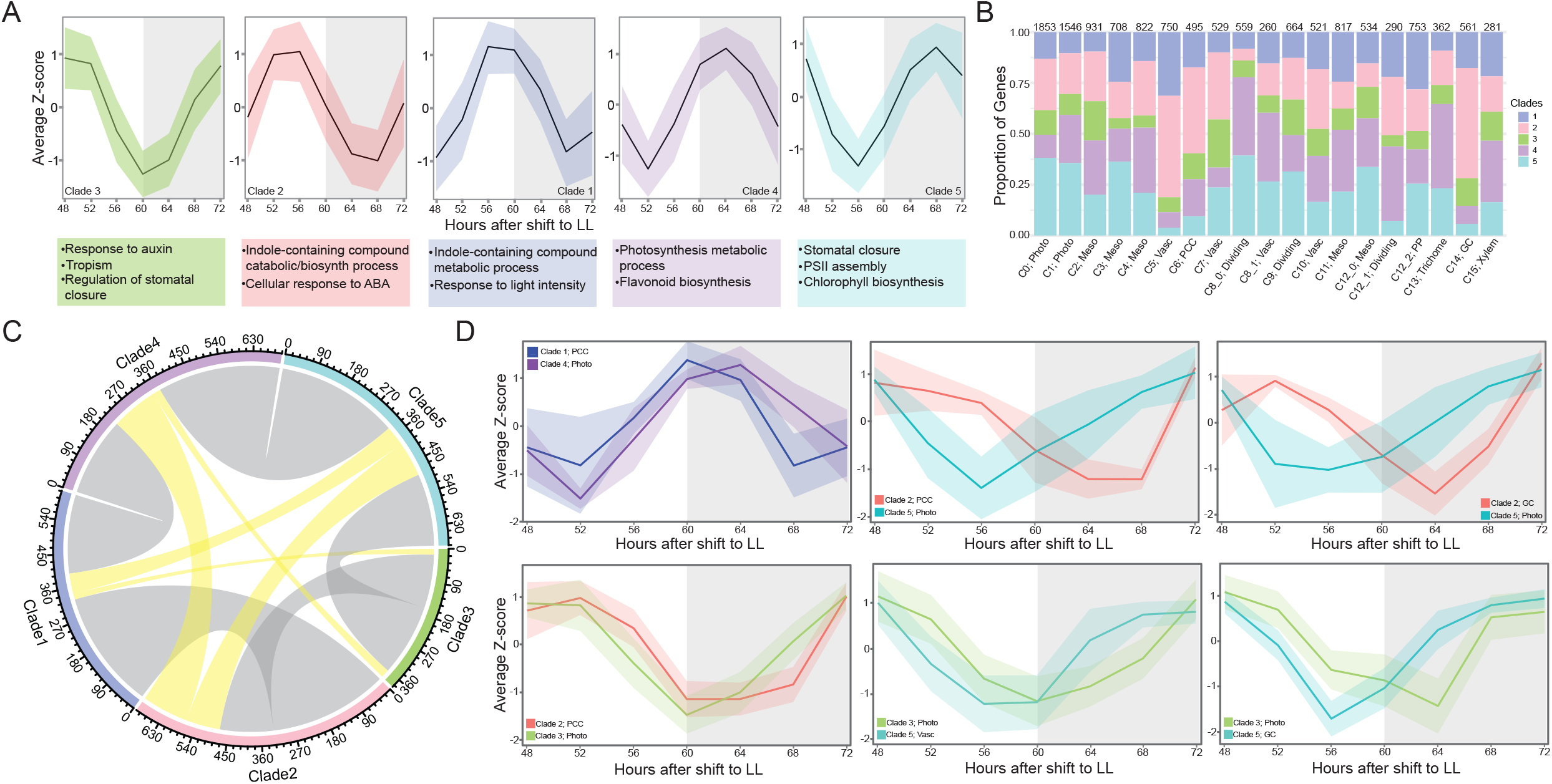
Temporal clades and cell-type-specific phase shifts. (A) WGCNA modules grouped into five clades, shown as mean z-score ± SD with representative GO enrichments. (B) Clade proportions by cluster. (C) Cross-cluster clade changes; ribbon width indicates gene number, and gray/yellow denote adjacent/nonadjacent shifts. (D) Mean z-score ± SD for gene sets with tissue-specific phase differences.

To analyze the biological processes enriched for the clades identified, we performed GO enrichment analysis (*Dataset S3*). The GO analysis identified some common and unique enrichment terms among the clades; the early morning clade (clade 3) had genes involved in auxin response, tropism and regulation of stomatal closure. Similarly, we saw enrichment of genes for indole-containing compound catabolic/biosynthesis processes and cellular response to ABA enriched in the midday clade (clade 2). The GO terms related to preparation for photosynthetic machinery were enriched in the second half of the day indicating the plant’s strategy to prepare for the energy harvesting processes of the next day (Fig. 3A).

To examine whether genes cycling across clusters exhibited similar phase patterns we identified genes that were found in multiple clades. The majority of genes (5611) were exclusive to one clade only, whereas 1540, 114, and 1 gene were found in 2, 3, and 4 clades respectively. For the genes present in two clades, we find most shifts occurred between adjacent clades (Fig. 3C; grey ribbons); however, a small fraction of genes exhibited more dramatic phase changes (Fig. 3C; yellow ribbons). Examining these genes at the cluster level revealed similar changes between clusters suggesting a common regulatory mechanism (Fig. 3D). Notably, we identified a handful of genes in our photosynthesizing cluster that showed phase changes when compared to phloem companion cell (PCC) or guard cell (GC), resulting in their segregation into different clades (Fig. 3D). As our analysis revealed that the same genes can have cell type specific expression patterns, we next aimed to construct cell type-specific gene regulatory networks to enhance our understanding of the unique regulatory mechanism at play in each cell type.

### Cell-type-resolved regulatory networks reveal shared and cell-type-specific targets of core clock regulators

We next constructed circadian single cell gene regulatory networks (scGRNs) to better characterize the ways in which circadian genes are differentially regulated across cell types. We implemented the GENIE3 algorithm (Huynh-Thu et al. 2010) on individual clusters while simultaneously limiting our input dataset to genes which are cycling in that specific cluster. In parallel, we iteratively shuffled each cluster’s data and ran GENIE3 to identify null distributions of each dataset (Huynh-Thu et al. 2010). The weight value corresponding to the 95th percentile was used as a weight cutoff for the unshuffled connections, leaving ∼156,000 TF-target connections spread across all clusters (Fig 4A, *Dataset S7*). Because networks were inferred independently within each cluster and restricted to genes classified as cycling in that cluster, absence of an edge should be interpreted cautiously and may reflect thresholding or cycling detection rather than true absence of regulation. Approximately 25% of the edges had TFs and targets assigned to the same WGCNA clade, while the remaining 75% were assigned to different WGCNA clades (Fig. 4B). Although GENIE3 does not assign regulatory sign, the comparison with the temporal co-expression clades provides a coarse view of whether predicted TF-target pairs tend to share or diverge in phase, which suggests we have captured instances of activation and repression.

**Figure 4.**
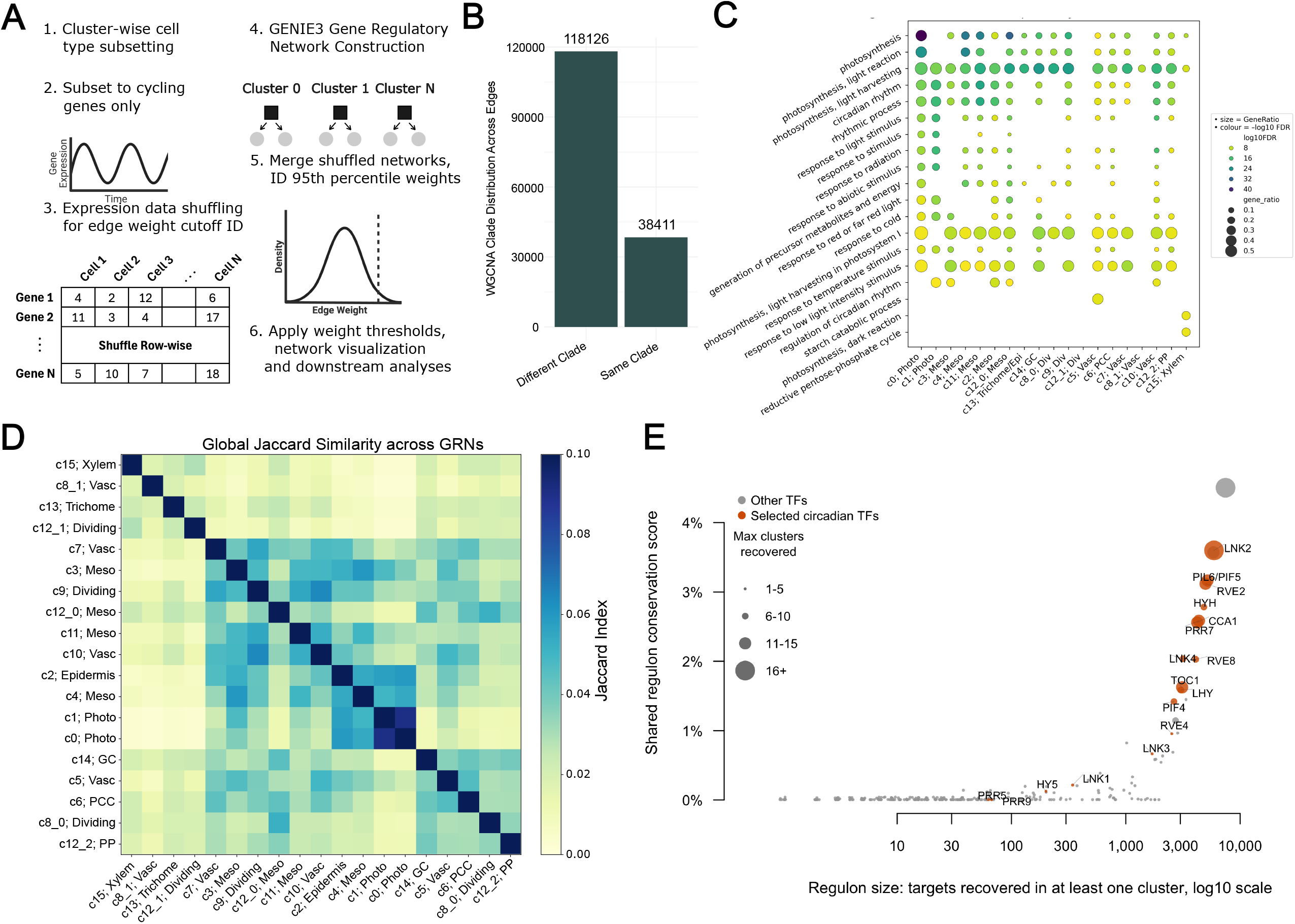
Cluster-resolved circadian single-cell gene regulatory networks (scGRNs). (A) Construction and filtering. (B) Edges linking TFs and targets in the same versus different WGCNA clades. (C) Top five GO biological-process enrichments per cluster. (D) Between-cluster Jaccard similarity of significant edges. (E) TF target-conservation scores; highlighted regulators are orange. Photo, photosynthetic; Meso, mesophyll; Epi, epidermis; GC, guard cell; Div, dividing; Vasc, vascular; PCC, phloem companion cell; PP, phloem parenchyma.

The scGRN target sets in individual clusters were broadly enriched for photosynthetic and circadian GO terms (Fig. 4C), reflective of the strong regulatory control the clock exerts on the timing of photosynthetic activity and other rhythmic processes (Hennessey and Field 1991; Harmer et al. 2000; Haydon et al. 2013; Dodd et al. 2014a). Enrichment was also identified for a number of environmental responsive terms, including temperature and light responses. Given that both light and temperature are key zeitgebers in plant circadian entrainment (Wang et al. 2022), examining the specific interactions underlying these enrichments could provide new insights into how clock entrainment is maintained in diverse cell types. Comparison of the scGRNs between clusters identified larger target overlaps between clusters of similar cell types (Fig. 4D), supporting that different cell types within a common tissue maintain some amount of conserved network structure. The patterns observed across our scGRNs were largely mirrored when comparing to scGRNs constructed from a recently published single nucleus circadian time course in Arabidopsis seedlings (Qin et al. 2025), particularly in mesophyll and guard cell clusters (Fig. S7, *Dataset S8*), where shared edges between clusters assigned to the same cell type often share 5-10% of TF-target edges between experiments, despite the dramatically different developmental processes taking place between whole seedlings and mature leaves (Kalve et al. 2014).

Network similarity was also observed at the individual TF level, where Jaccard index based hierarchical clustering highlighted similarities within cell types, including for CCA1, LNK2, TOC1, and PRR7 (Fig. S6). LNK2 in particular showed high conservation between many clusters, consistent with its role as a general circadian co-activator. We reasoned that some of these conserved connections between pairs of scGRNs might be more broadly maintained across more than two clusters. To explore this, we calculated shared regulon conservation scores for each TF, identifying TFs with larger fractions of targets that recurred across multiple cluster-resolved scGRNs (Fig. 4E). This analysis identified many clock TFs, such as CCA1, PRR7, and LNK2, predicting a substantial amount of conserved function across different cell types for these circadian regulators. It should be noted that lower conservation metrics may stem from technical limitations of nuclei preparation or cycling cutoffs which is why we focus on the high ranked TFs. Based on the high relative conservation score of CCA1, we examined the shared targets between pairs of scGRNs and found that the similarity between CCA1-centered networks was largely driven by which and how many photosynthesis-associated targets were identified (Fig. S8). A suite of 11 light-harvesting chlorophyll a/b-binding proteins (*LHCB*) with CCA1 edges spanned at least 3 clusters, whereas 7 of the 9 CCA1 edges found in only one or two clusters were associated with Photosystem II repair, with 6 of these 20 CCA1-target pairs (Fig. S9A) supported by previously published ChIP-seq data (Nagel et al. 2015). The combined action of broad light harvesting influence and cell type specific PSII repair targets of CCA1 highlight the complex and layered regulation by the clock to fine-tune photosynthetic activity.

In addition to the similarity of photosynthesis-associated targets across clusters, we also noted that auxin responsive genes in CCA1-centered networks contained a high proportion of edges restricted to a single cluster (Fig. S9B). More than 20% of all genes assigned to auxin-responsive GO categories were identified in CCA1-centered networks in at least one cluster, with edges called between CCA1 and 64 unique targets. A majority of these CCA1-target pairs were identified in Photosynthetic clusters c0 and c1; however, 13 pairs were unique to clusters other than c0 or c1, and an additional 21 pairs were present in at least one non-photosynthetic cluster. To further support these predicted TF-target connections, we again leveraged previously published CCA1 ChIP-seq experiments (Nagel et al. 2015), identifying a subset of 18 TF-target edges supported by bulk ChIP-seq binding (Fig. S9, S10A). While many of these ChIP-validated edges were only found in Photosynthetic cells, more than half were also present in at least one other tissue or cell type. These two regulons highlight instances where CCA1 is predicted to broadly target light-harvesting genes, whereas there appears to be greater cell-type restriction for auxin targets.

### CCA1 and LNK2 target modules reveal shared clock-associated programs with distinct cell-type contexts

We next focused on CCA1 and LNK2 because they ranked among the most highly conserved clock associated regulons (Fig. 4E), yet their detected target sets differed in size and cell-type distribution. This comparison allowed us to ask whether CCA1, a canonical DNA-binding clock TF, and LNK2, a clock-associated transcriptional co-regulator (Xie et al. 2014), converge on common target programs or implement distinct cell-type contexts. To resolve the conserved and cell-type specific regulation and reduce cluster-level sparsity while retaining major tissue context, we merged clusters into five major cell types: Photosynthesizing cells, Mesophyll, Epidermis, Vascular and Dividing. Notably, the c0 cluster was removed from the merged Photosynthesizing cells due to moderately higher ambient background noise that led to spurious co-association of many cell types, enabling us to more confidently associate predicted regulation with actively photosynthesizing cells. We next performed GO enrichment analysis on the TF targets for each merged cluster and compared the significant terms. There were 144 GO terms shared between CCA1 and LNK2, with an additional 129 terms unique to LNK2 and 24 unique to CCA1. Interestingly, we found that 90% of CCA1 and 62% of LNK2 predicted targets overlapped (Fig. 5A, purple nodes) suggesting that a substantial fraction of the detected CCA1 regulon is embedded within broader LNK2-associated target programs, while LNK2 also contributes additional predicted targets not recovered for CCA1 under our filtering criteria.

**Figure 5.**
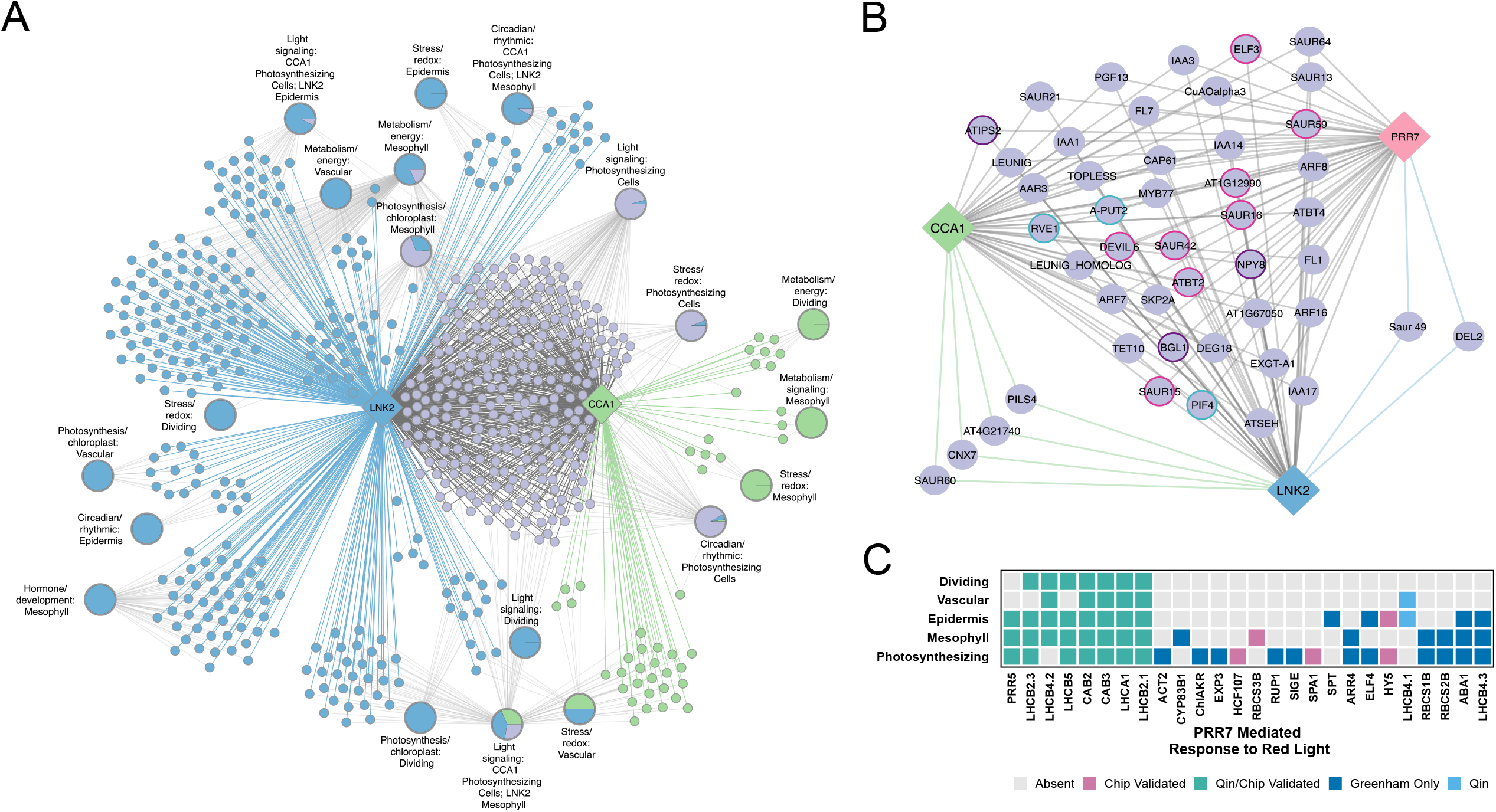
Clock TF coregulatory networks. (A) CCA1/LNK2 targets linked to GO supertheme/cell-context modules. Green, blue, and gray denote CCA1-only, LNK2-only, and shared targets and interactions; pies show module target composition. (B) Auxin-response network; gray, green, and blue edges denote CCA1/LNK2/PRR7, CCA1/LNK2, and PRR7/LNK2 regulation. (C) Heatmap of significant PRR7-centered response-to-red-light (GO:0010114) edges.

Further examination of the CCA1 and LNK2 targets revealed many shared enriched biological pathways, especially for circadian regulation, light signaling, and photosynthesis-related processes; however, the cell-type context differed in many of these contexts. For example, light-signaling-associated targets were detected in LNK2-enriched epidermal modules and in CCA1-associated photosynthesizing-cell modules, suggesting that similar clock-linked processes may be represented in distinct cell-type contexts (Fig. 5A, *Dataset S9*). More broadly, LNK2-associated modules spanned multiple biological themes and cell contexts, including stress/redox responses, metabolism/energy, and hormone- or development-associated processes (Fig. 5A, blue nodes), whereas CCA1-enriched modules were detected in a smaller set of stress/redox and metabolism-associated contexts (Fig. 5A, green nodes). Shared CCA1/LNK2 targets were distributed across several modules, including circadian/rhythmic, light signaling, and metabolism-associated processes, indicating that overlap between these regulators is not confined to a single pathway, but instead spans multiple clock-associated outputs (Fig. 5A, purple nodes). Because module assignment depends on cycling-gene inclusion and scGRN edge threshold, these patterns are best interpreted as candidate cell-type-resolved regulatory programs rather than exhaustive TF-specific regulons.

Among the co-regulated pathways, we noted that 45 genes in the enriched response to auxin term (GO:0009733) were predicted targets of both CCA1 and LNK2 in photosynthesizing cells, suggesting potential co-regulation of the auxin response in this cell type (Fig. 5B, *Dataset S10*). Additionally, LNK2 has two predicted targets that do not overlap with CCA1 predicted targets. As LNK2 does not have a DNA binding domain and functions as a co-activator (Rugnone et al. 2013), we wondered whether it might be co-regulating these genes with another clock TF. We found that PRR7 targets were also enriched for the auxin response GO term, with 41 predicted targets shared by all three TFs (Fig. 5B, grey edges), four predicted targets regulated by CCA1/LNK2, and two unique to PPR7/LNK2. The shared co-regulation architecture is further supported by the high target overlap among the TFs as PRR7 shares 93% of its targets with LNK2, while CCA1 and PRR7 share ∼75% of their targets. These overlapping predicted target sets place PRR7 as an additional candidate clock regulator within this auxin-associated module.

Expanding beyond the CCA1/LNK2 comparison, PRR7-centered networks further highlighted light perception and photosynthetic capacity as recurring predicted targets of core clock-associated regulators. Specifically, we identified many edges associated with red light signaling. Many of these PRR7-*LHC* edges were supported by previously published ChIP-seq binding data (Liu et al. 2016) and the Qin et al. single cell data set (Fig. S10) (Qin et al. 2025), again highlighting the strong connection between the circadian clock and the daily timing of photosynthetic capacity. Among edges detected, we observed several well-described light signaling components, including targets implicated as PRR7 bound in the referenced ChIP-seq dataset(Liu et al. 2016), such as *ELONGATED HYPOCOTYL5 (HY5*) and *SUPPRESSOR OF PHYA-105 1 (SPA1*) (Fig. 5C). Most of the edges identified in different cell types were associated with light harvest and capture, again highlighting the fundamental role of the clock in properly timing photosynthetic capacity. Ultimately, these findings further strengthen our understanding of the role of the plant circadian clock, shifting the paradigm from a uniform global oscillator to a highly cell type specific regulatory network. Together, these predicted networks support a model in which core clock-associated regulators contribute to both distributed programs, such as photosynthetic light capture, and more cell-type-restricted programs, such as auxin-associated growth responses.

### Spatial transcriptomics confirms tissue organization and reveals spatial heterogeneity in clock gene expression

To recover the spatial information lost during nuclei isolation, we performed Xenium spatial transcriptomic profiling on formalin-fixed, paraffin-embedded sections of mature Arabidopsis leaves at two circadian time points, capturing dawn (ZT48, Fig. 6A) and dusk (ZT60, Fig. 6B) spatial gene expression information across a 100-gene panel (*Dataset S11*). Leaves were sliced in a paradermal orientation in order to preserve as much spatial context as possible, leading to mesophyll-dominated tissue sections that contain vascular bundles of variable lengths across the leaf and guard cell studded epidermis surrounding the ground tissue. Independent clustering of Xenium transcript profiles following Baysor-guided cell segmentation (Petukhov et al. 2022) and minimal manual curation identified six clusters in each time point, corresponding to one guard cell cluster and varying numbers of epidermal, vascular, and mesophyll cell type populations (Fig 6C). Cluster-resolved expression of cell type specific markers, including epidermal (*RESPONSE TO DESICCATION 22*; *RD22*), vascular (cytochrome P450, family 83, subfamily B, polypeptide 1; *CYP83B1*), mesophyll (*CHLOROPHYLL A/B BINDING PROTEIN 1*; *CAB1*), and guard cell (*FMA*) populations, allowed us to directly associated clusters with specific cell types (Fig. 6A-B, *Dataset S12, S13*). Expression across the 100-gene panel strongly supported the specific cell type assignments for each of these clusters while also cataloguing expression of genes involved in the clock, light responses, photosynthesis, stress, hormone responses, specialized metabolism, nutrient transport, and cell wall organization (Fig 6D).

**Figure 6.**
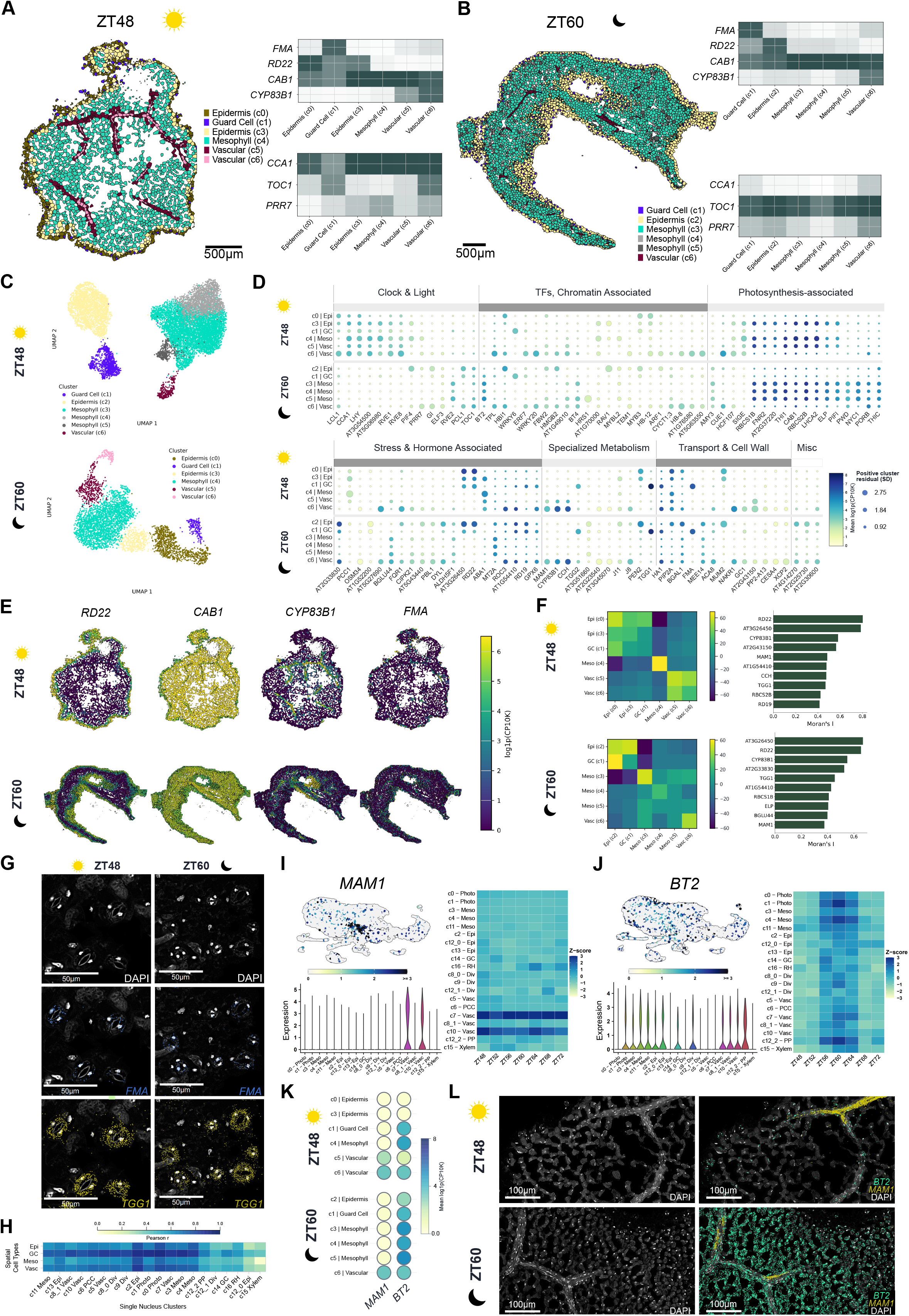
Xenium profiling of leaf organization and temporal expression. (A–B) ZT48/ZT60 spatial clusters and cluster-level marker/clock heatmaps. (C) Spatial-cluster UMAPs. (D) The 100-gene panel; color indicates mean log1p counts per 10,000 (CP10K) and size the positive cluster residual. (E) Spatial *RD22, CAB1, CYP83B1,* and *FMA* expression. (F) Cluster-neighborhood enrichment and the ten highest-Moran’s-I genes per time point. (G) DAPI, *FMA,* and *TGG1* maps. (H) Pearson correlations of ZT60–ZT48 differences between matched Xenium cell types and single-nucleus clusters. (I–J) Single-nucleus UMAP, violin, and temporal z-score plots for *MAM1* and *BT2*. (K) Mean Xenium-cluster *MAM1* and *BT2* expression. (L) DAPI, *BT2,* and *MAM1* maps. DAPI is gray; FMA/TGG1 are blue/yellow, and BT2/MAM1 cyan/yellow. Sun/moon denote ZT48/ZT60. Scale bars, 500 µm (A–B), 50 µm (G), and 100 µm (L). Photo, photosynthetic; Meso, mesophyll; Epi, epidermis; GC, guard cell; RH, root hair; Div, dividing; Vasc, vascular; PCC, phloem companion cell; PP, phloem parenchyma.

Direct spatial mapping of the marker genes further reinforced these cell type assignments. *RD22* was concentrated in epidermal regions, *CAB1* was broadly distributed throughout the mesophyll, *CYP83B1* followed vascular bundles, and *FMA* marked a scattered cell type matching guard cell morphology (Fig 6E). Neighborhood enrichment analyses similarly recovered the expected spatial organization of the paradermal leaf section. Epidermal and vascular populations preferentially occupied shared neighborhoods, while guard cells were depleted from other guard cell neighborhoods and associated more frequently with the epidermis, consistent with previously described stomatal spacing (Fig 6F)(Nadeau and Sack 2002; Hara et al. 2007). Gene-level Moran’s I analysis, a measure used to evaluate spatial autocorrelation, further identified strongly structured, nonrandom expression patterns. Cell type-associated genes, including *RD22* and *CYP83B1*, were among the most spatially autocorrelated genes at both time points, with *THIOGLUCOSIDE GLUCOHYDROLASE 1 (TGG1*) and *METHYLTHIOALKYLMALATE SYNTHASE 1* (*MAM1*) also ranking among the highest scoring genes at each time point (Fig 6F). To confirm these patterns independently of cell segmentation and cluster averaging, we examined individual transcript calls at high spatial resolution in the Xenium Explorer. *TGG1* transcripts were concentrated around morphologically distinct and *FMA*-expression guard cells at both ZT48 and ZT60 (Fig 6G), supporting guard cell enrichment of *TGG1* expression.

We next asked whether the observed circadian differences detected in intact tissues were consistent with the single nucleus time course. We collapsed similar Xenium cell types at each time point to calculate the difference in transcript accumulation between the two time points and used Pearson correlations to compare these changes with the corresponding difference across the single nucleus time course (Fig 6H). Correlations across all genes in the Xenium panel revealed broad agreement between the two datasets, indicating that genes with relatively higher expression at ZT48 or ZT60 in the spatial dataset generally displayed concordant time-point differences in the single nucleus dataset. This broad cross-platform agreement supports the reproducibility of cell type resolved temporal expression patterns across the spatial and single nucleus datasets.

At the level of individual genes, we focused on *MAM1* and *BTB AND TAZ DOMAIN PROTEIN 2 (BT2*) to further explore concordance of the spatial and single nucleus experiments. *MAM1* expression was largely restricted to c7 and c10, a subset of vascular clusters, but did not vary significantly across the single nucleus time course (Fig 6I). In contrast, *BT2* was detected across most clusters, but displayed a strong rhythm peaking around ZT60 in the single nucleus time course (Fig 6J). Xenium cluster averages captured these differences, where *MAM1* showed strong accumulation in the vasculature at both ZT48 and ZT60, and *BT2* showed higher accumulation across all cell types at ZT60 compared to ZT48 (Fig 6K). Interestingly, modest accumulation of *BT2* was observed in a subset of spatial clusters at ZT48 and could correspond to the handful of clusters in the single nucleus data with relatively higher expression at ZT48 (Fig 6J, c11 mesophyll and c12_2 phloem parenchyma). To better understand these patterns, we directly examined transcript localization for *MAM1* and *BT2* in the Xenium explorer (Fig 6L). We observed spatially distinct vascular cell populations with high *MAM1* expression across both time points, supporting our observation of expression in only a subset of vascular cell types in the single nucleus time course. *BT2* expression was broad and high at ZT60 but was primarily observed in guard cells and vasculature at ZT48. At both time points, *BT2* expression was more broadly distributed through the vasculature than the accumulation pattern observed for *MAM1*.

We next examined the spatial organization of core clock transcripts across the two circadian time points (*Dataset S14*). Across the full tissue sections, transcript abundance followed the expected time-of-day patterns, with high and low *CCA1* expression at ZT48 and ZT60, respectively, and the reciprocal pattern observed for *TOC1* (Fig 7A). Cell type-resolved quantification confirmed that *CCA1* was higher at ZT48 and *TOC1* was higher at ZT60 in epidermal, guard cell, mesophyll, and vascular cell types (Fig 7B); however, the magnitude of these differences was not uniform across cell types. *CCA1* was relatively lower in abundance in guard cells and a subset of mesophyll cells at ZT48 and had relatively higher abundance in guard cells and vasculature at ZT60. *TOC1* had relatively higher abundance in guard cells and vasculature at ZT48, and relatively lower abundance in a subset of mesophyll cells at ZT60. We observed similar patterns in other clock components, where clock genes at the ZT corresponding to their lowest global expression tended to have increased accumulation in vascular and guard cells (Fig. S11).

**Figure 7.**
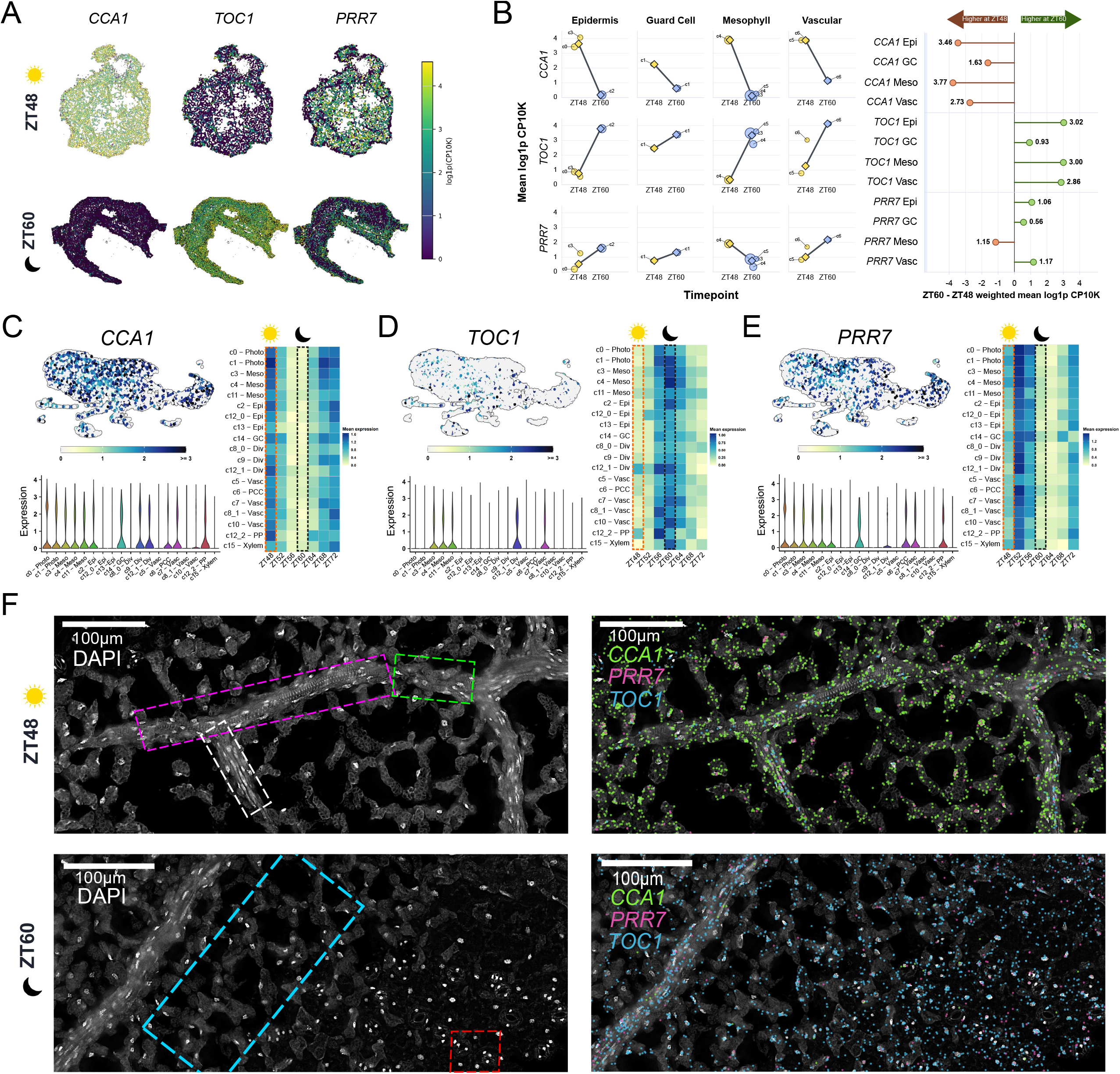
Spatial variation in core clock expression. (A) Spatial CCA1, TOC1, and PRR7 expression at ZT48 and ZT60. (B) Expression by cell type. Points denote spatial clusters and connected diamonds cell-type-weighted means; the summary shows ZT60–ZT48 mean log1p CP10K differences (brown, ZT48 higher; green, ZT60 higher). (C–E) Single-nucleus UMAP, violin, and temporal-expression plots for CCA1, TOC1, and PRR7. (F) DAPI (gray), CCA1 (green), TOC1 (blue), and PRR7 (magenta). Dashed outlines mark ZT48 vascular regions (white, pink, green), ZT60 mesophyll (blue), and ZT60 guard cells (red). Sun/moon denote ZT48/ZT60. Scale bars, 100 µm.

Much more complex spatial distributions were identified for *PRR7*, whose peak expression falls between the two spatial time points (Fig 7A). Unlike *CCA1* or *TOC1*, which changed in the same direction across all four broad cell types, the direction of the *PRR7* difference varied among tissues. *PRR7* expression was higher at ZT48 in mesophyll cells, but was higher at ZT60 in epidermal, guard cell, and vascular cell types (Fig 7B). Although two spatial time points cannot distinguish changes in phase from differences in amplitude or waveform, these results demonstrate that temporal changes in *PRR7* transcript abundance are not uniform across cell types.

These high-resolution insights into the spatial organization of clock TF accumulation drove us to return to our single nucleus data to re-examine *CCA1, TOC1,* and *PRR7* (Fig 7C-E). *CCA1* mean expression across the single nucleus clusters at ZT48 and ZT60 was fairly uniform, and did not seem to capture the relative guard cell depletion at ZT48 or the relative vascular enrichment at ZT60 observed in the spatial data, though *CCA1* expression was increased by ZT64 in a subset of vascular clusters (Fig 7C). *TOC1* mean expression was far less uniform across the single nucleus clusters at both time points, with notably high expression in a subset of phloem parenchyma at ZT48 and low expression in a subset of mesophyll clusters at ZT60 (Fig 7D). *PRR7* mean expression was variable across clusters at both time points, with moderately more accumulation in several photosynthetic and mesophyll clusters, and notable accumulation in phloem companion cells and guard cells at ZT60 (Fig 7E). The patterns and distributions of expression for *TOC1* and *PRR7* in the single nucleus time course largely supported those observed in cluster-resolved averages in the spatial datasets, while the observed *CCA1* spatial accumulation was not fully captured by the nuclear profiles. These differences can likely be attributed at least in part due to the experimental approaches, where Xenium spatial transcriptomics primarily capture mature, accumulated mRNA and single nucleus-based transcriptomics primarily capture actively or recently transcribed mRNA present only in the nucleus.

Finally, we examined individual transcript localization for *CCA1, TOC1,* and *PRR7* through the Xenium Explorer to assess these spatial distributions at higher resolution. *CCA1* transcript localization across the two circadian timepoints largely captured the cell-type-resolved differences discussed above, including increased *CCA1* expression in vascular and guard cells at ZT60 (Fig. 7F, GCs outlined in red region). *TOC1* transcripts were primarily observed in a subset of vascular cells at ZT48 (Fig 7F, magenta and green outlined regions) and were visibly reduced in a subset of mesophyll cells at ZT60 (Fig 7F, blue outlined region). *PRR7* transcripts at ZT48 were present across the mesophyll, as well as in a subset of vascular cells (Fig. 7F, white and pink outlined regions) but were notably absent in the mesophyll surrounding the vasculature at ZT60 (Fig 7F, blue outlined region). These expression patterns support prior observations that distinct cell types display altered circadian rhythms and provide excellent spatial resolution to understand circadian expression in the mature Arabidopsis leaf. When considered alongside the complete single nucleus time course, these spatially resolved differences support the conclusion that broadly shared circadian expression programs are modulated by cell type context within the mature Arabidopsis leaf.

## Discussion

Unlike animals, plants lack a central nervous system, raising the question of how autonomous local clocks are coordinated across tissues. Here, we combined a 24-h mature-leaf snRNA-seq time course and spatial transcriptomics along with cell-type-resolved scGRNs to examine how circadian regulation varies across leaf cell types and tissue contexts. Because many of the common marker genes used for identification (Wang et al. 2021; Xia et al. 2022; Lee et al. 2023) are themselves rhythmic (Romanowski et al. 2020) (Fig. 1B), we masked rhythmic genes during integration to recover cell-type identity. We resolved 20 clusters spanning the major mature-leaf cell types, which were grouped into broad mesophyll (M), epidermal (E), dividing (D), vascular (V), and root-hair-associated (RH) populations (Fig. 1D-F). Xenium profiling at ZT48 and ZT60 provided orthogonal support for these annotations while preserving positional information, which was particularly important in the leaf, where closely related photosynthetic and mesophyll clusters may represent transcriptional states that are difficult to parse without spatial context.

Our pseudo-bulk analysis showed cycling of *CCA1*, *LHY*, *PRRs*, *RVEs*, *LNKs* and other clock genes (Fig. 2A) with peak expression at their expected time of the day (Hsu and Harmer 2014). Our cycling analysis of individual clusters revealed that many of these core clock genes cycle in many clusters (Fig. 2B), supporting the presence of functional oscillators across major mature-leaf cell types; however, several clock components, including *PRR5* and *LUX*, cycled only in subsets of clusters, suggesting cell-type differences in clock output or detectability. The spatial data further supported our conclusion that circadian gene expression is broadly distributed throughout the leaf (Fig. 6D) and confirmed that time-of-day differences in clock transcript accumulation are detectable within intact leaf domains, anatomically supporting the cluster-resolved rhythms. Together, these results are consistent with a model in which the mature leaf contains multiple local circadian oscillators operating within a shared tissue framework, where cell identity and spatial context influence the phase, amplitude, and downstream regulatory outputs of clock-associated gene expression.

To validate the temporal structure of cluster-resolved rhythmic genes, we used WGCNA (Langfelder and Horvath 2008) to group cluster-resolved rhythmic genes into five temporal clades corresponding to expected morning, midday, evening, and night expression programs (Fig. 3A). These clades recovered canonical clock phase relationships and were enriched for known circadian-regulated processes, including auxin response (Covington and Harmer 2007; Xue et al. 2020), ABA/stomatal regulation (Hubbard and Webb 2015), photosynthesis (Hennessey and Field 1991; Harmer et al. 2000; Booij-James et al. 2002; Edwards et al. 2006; Haydon et al. 2013; Dodd et al. 2014a; Dodd et al. 2014b) and defense-associated pathways (Melotto et al. 2006; Goodspeed et al. 2012; Zheng et al. 2015). Early morning clades were enriched for auxin responses, coinciding with previous findings that auxin responses are clock gated to increase from dusk to dawn (Covington and Harmer 2007; Xue et al. 2020). We also identified enrichment of salicylic acid (SA) signalling pathways in our morning clade, in accordance with when plants are highly vulnerable to pathogen infection (Melotto et al. 2006; Goodspeed et al. 2012; Zheng et al. 2015). Importantly, genes assigned to different clades across clusters revealed cases in which the same gene exhibits altered phase relationships among cell types, supporting cell-type-specific tuning of circadian outputs (Fig. 3C).

Cluster-resolved scGRNs built from cycling genes provided a hypothesis-generating framework for identifying clock-associated regulatory modules that are broadly shared or preferentially detected in specific cell-type contexts (Fig. 4A) (Nagel et al. 2015; Liu et al. 2016) nevertheless supports the biological relevance of the inferred networks and provides a basis for exploring how clock-associated regulators are integrated across cell types (Fig. S10). Interestingly, when comparing genes that were regulated across multiple clusters to genes regulated only in a single cluster, we identified broad predicted regulation by CCA1 on light harvesting components, compared to more limited regulation by CCA1 on photosystem II repair components. One possible explanation is that the distinct mesophyll clusters we have identified are being clustered at least in part by their photosynthetic status; however, GO enrichment on cluster biomarker sets identified a significant enrichment for PSII repair terms in only one of five mesophyll clusters (Cluster 1, *Dataset S3*).

Direct links tying the circadian clock to steady state PSII repair cycles are largely missing; however, endogenous cycling of phosphorylation status of repair components has been documented (Booij-James et al. 2002; Edwards et al. 2006; Dodd et al. 2014b). Additionally, clock gating of PSII repair components has been described in response to stress conditions, including gating of *SIGMA FACTOR5 (SIG5)*, a key regulator of the psbDC operon that is involved in D protein cycling (Edelman and Mattoo 2008; Noordally et al. 2013; Cano-Ramirez et al. 2023). The nuclear encoded protease *DEGRADATION OF PERIPLASMIC PROTEINS 8 (DEG8)* which is also involved in D protein cycling, was identified as a CCA1 target through scGRN predictions, and is supported by previously published CCA1 ChIP-seq binding (McClure et al. 1989; Nagel et al. 2015). Further examination of the relationship between CCA1 and DEG8 could solidify a regulatory link between the clock and steady state PSII repair cycles.

In addition to photosynthetic regulation, the clock gates auxin biosynthesis, transport, and sensitivity (Jouve et al. 1999; Covington and Harmer 2007), but how this regulation varies among mature-leaf cell types remains unclear. A recent study examined circadian gene expression in Arabidopsis seedlings at the single nucleus resolution, identifying a subset of six *SAUR* genes cycling exclusively in the shoot epidermis (Qin et al. 2025). Our scGRN-based approach identified 64 auxin-associated CCA1 targets across all clusters, including a subset of 21 genes previously identified as potential CCA1 targets through bulk ChIP-seq. Many of these genes robustly cycle in multiple clusters, but CCA1-target edges were recovered only in a subset, suggesting candidate cell-type-restricted regulatory relationships that would be missed in bulk analyses. One example matching these criteria is *SMALL AUXIN UPREGULATED 15* (*SAUR15*), a ChIP-seq validated CCA1 target (McClure et al. 1989; Nagel et al. 2015) that is predicted to be CCA1 regulated in Photosynthetic c1, but not in Vascular c5 or Phloem Companion Cells c6. These patterns suggest that clock regulation of auxin-associated genes is not uniform across the leaf and may contribute to cell-type-specific growth or signaling outputs.

PRR7-centered networks further linked the clock to light signaling and photosynthetic capacity (Fig 5B) (Kaczorowski and Quail 2003; Ito et al. 2007; Liu et al. 2016), where predicted PRR7 targets included light-responsive regulators and LHC genes across multiple cell types (Fig. 5C).These included key photomorphogenic regulators such as *HY5* and *SPA1*, which are central to red/far-red light signaling and cell elongation in the hypocotyl (Saijo et al. 2003; Hoecker 2017). PRR7 connections to *HY5* were identified in both mesophyll and epidermal clusters, while *SPA1* connections were exclusively found in mesophyll clusters. The presence of PRR7–*LHC* (light-harvesting chlorophyll complex) edges in multiple cell types further suggest that PRR7 may contribute to both broadly distributed light-harvesting programs and more cell-type-specific light-signaling outputs in the mature leaf.

While it is well established that clock transcription factors co-regulate distinct clock outputs (Li et al. 2025), Night Light-Inducible and Clock Regulated proteins (LNKs) are specifically recognized for their co-activation activity. Because LNKs lack a native DNA-binding domain, they recruit and interact with morning-phase TFs, such as CCA1, LHY, RVE4, and RVE8, to modulate gene expression (Xie et al. 2014). LNK2-associated modules overlapped extensively with CCA1- and PRR7-associated programs, including rhythmic, light-responsive, photosynthetic, metabolic, and hormone-related processes, but often in distinct cell-type contexts (Fig. 5). These predictions suggest that LNK2 marks co-regulator-associated clock output programs, although future perturbation or spatially resolved validation will be required to distinguish direct recruitment from indirect association.

Although snRNA-seq revealed broad cycling of clock components, subtle differences in low-abundance transcripts or small cell populations were difficult to interpret from cluster averages alone (Fig 7C-E) Xenium spatial transcriptomics preserved tissue organization (Fig 6A-D) and confirmed many cell-type-resolved temporal differences while revealing localized patterns within vascular, mesophyll, epidermal, and guard cell populations (Fig 6H). Differences between platforms, such as those observed for CCA1, may reflect mRNA stability, compartmentalization, detection chemistry, or resolution. Thus, the spatial data both validate and extend the snRNA-seq time course by resolving local patterns obscured by dissociation-based clustering.

*CCA1* and *TOC1*, which peak near ZT48 and ZT60, respectively, were both present in higher abundance in vasculature and guard cells when at their global minimum. Previous studies have reported that vascular clocks maintain more robust rhythms than mesophyll clocks and that guard cells can exhibit distinct circadian periods relative to adjacent tissues (James et al. 2008; Yakir et al. 2011; Endo et al. 2014). Our observations are consistent with differences in clock timing or transcript accumulation across these cell types; however, it is not possible to determine from two time points whether these patterns reflect differences in phase, period, amplitude, waveform, or transcript turnover. *PRR7* provided a distinct example, with higher abundance in mesophyll cells at ZT48 and higher abundance in epidermal, guard cell, and vascular populations at ZT60. Because *PRR7* peaks between the sampled time points, this pattern is consistent with an earlier rise and decline in mesophyll relative to other tissues, although differences in amplitude or waveform could produce a similar result. Together, these data indicate that temporal changes in clock transcript abundance are not uniform across the mature leaf.

Prior work has described clock waves across leaves and vein-associated phase relationships (Fukuda et al. 2007; Wenden et al. 2012), and our spatial data extend this view by identifying cell-type- and region-specific differences in clock transcript accumulation. Future spatial time courses across multiple cycles, combined with live reporters and cell-type-specific perturbations, will be needed to distinguish phase, period, amplitude, waveform, and transcript-turnover effects. More broadly, our findings show that cluster-resolved snRNA-seq captures only part of the spatial regulatory landscape and establish a framework for testing how tissue organization shapes circadian regulation across the mature Arabidopsis leaf

## Material and Methods

### Plant Growth and Sampling

Arabidopsis Col-0 seeds were surface sterilized for four hours by vapor phase sterilization (Clough and Bent, 1998). Sterilized seeds were imbibed with water. Seeds were plated on half strength Murashige Skoog (MS) agar with 1% sucrose and a pH of 5.7 buffered with Potassium Hydroxide (KOH). Seeds were then stratified at 4°C for two days in complete darkness. Following stratification plates were placed in a chamber with 12L/12D photoperiod (light intensity 130 μmol s^−1^) at 22°C. Seedlings were entrained to 12L/12D conditions for 11 days. On the 11th day, starting at subjective dawn, seedlings were grown in constant light for 48 hours before the start of the time course. Leaf samples of 180-250 seedlings were harvested by cutting across the root-shoot junction in 2 replicates for single-nuclei RNA-seq every 4 hours over a 24-hour period starting at the subjective dawn of the 13^th^ day.

### Nuclei Isolation

Leaf tissue was collected in a petri plate containing 5 ml of pre-chilled NIB (10mM Tris-HCL-pH 7.4, 10 mM NaCl, 3 mM MgCl, 0.5mM Spermidine, 0.2 mM Spermine, 0.5% Bovine Serum Albumin with 1:1000 Roche Complete Protease Inhibitor and 0.03 U/μl of RNase inhibitor but without detergents). Leaf samples were chopped at 4°C for 3 min on a petri plate containing 5 ml of NIB, with a razor blade. Finely chopped leaf samples were then shifted to 50 ml falcon tubes, ∼ 15 ml of NIB was added to make the final volume of sample 20 ml. Samples were incubated on a rotating platform for 5 min to facilitate cell release. After the first incubation, samples were filtered via 70 μm and 40 μm cell strainers to remove leaf debris. Triton X-100 and NP-40 were added to final concentration of 0.05%, and extracts were incubated on a rotating platform at 4°C for 10 min. The nuclei suspension was then centrifuged at 500 x g for 5 min at 4°C. The nuclei pellet was then resuspended in 5 ml of NIB without detergents and was again pelleted at 500 x g for 5 min at 4°C. The pellet was resuspended in 750 μl of resuspension buffer (1X PBS + 1% BSA + 0.03 U/ul RNaseI). Nuclei suspension was then shifted to the Nuclei Isolation column and collection tube of the Chromium nuclei isolation kit (PN-10000494, 10X Genomics, Pleasanton CA). Samples were then centrifuged at 16000 x g for 40 seconds. Pelleted nuclei were finally resuspended in 500 μl of resuspension buffer. Nuclei were counted manually with a hemocytometer. The 32k nuclei were loaded onto the 10x Genomics Chip M, which was then processed with 10x Genomics Chromium X controller, and sequencing libraries were constructed following manufacturer’s instructions except for cDNA amplification step where 13 cycles were used i.e. 2 additional cycles to prepare cDNA from nuclei (PN-1000268 number for the kit, 10X Genomics, Pleasanton CA). Total time from leaf harvest to loading was 30-40 min.

### Xenium spatial transcriptomics tissue collection

Plants used for Xenium spatial transcriptomics were grown under the same entrainment conditions described above. Briefly, Arabidopsis thaliana Col-0 seedlings were grown under a 12 h light/12 h dark photoperiod at 22°C, transferred to constant light, and collected at the indicated circadian time points. For spatial transcriptomic profiling, leaf tissue was collected at ZT48 and ZT60, corresponding to subjective dawn and subjective dusk, respectively. For each time point, one biological replicate was collected, consisting of two to three sections. Tissue was collected as quickly as possible and immediately transferred into a fresh fixative to preserve RNA integrity.

### Formalin fixation, paraffin embedding, and sectioning

Fixation and paraffin embedding of Arabidopsis leaves was adapted from previous work(Sheat et al. 2020). Briefly, leaf tissue was fixed in fresh 10% neutral-buffered formalin (Sigma-Aldrich, HT501128-4L) using vacuum infiltration. Tissue was transferred into a 50 mL syringe containing fixative, the syringe tip was sealed, and vacuum pressure was applied in repeated ∼30 second intervals until the tissue appeared fully infiltrated. Samples were maintained in formalin for 2 hours at room temperature. Following fixation, tissue was washed twice in 1× PBS for 15 min each at room temperature. Samples were then dehydrated through an ethanol series consisting of 30%, 50%, and 70% ethanol for 30 min each at room temperature, followed by storage at 4°C until further processing. Dehydration was completed with 90% and 100% ethanol for 30 min each.

Dehydrated samples were cleared through ethanol:xylene mixtures of 2:1, 1:1, and 1:2 for 45 min each at room temperature, followed by 100% xylene for 45 min. Tissue was infiltrated with xylene:paraffin mixtures of 2:1, 1:1, and 1:2 for 1 h each at 60°C, followed by incubation in pure molten paraffin for 1 h at 60°C. Samples were embedded in Paraffin Wax (Sigma-Aldrich, Cat #327212) using Flat Bottom Embedding Capsules and oriented to obtain paradermal sections. Paraffin blocks were cooled, mounted onto tissue cassettes, and stored at 4°C until sectioning.

Paraffin sections were cut at 10 μm using a Reichert-Jung 820-II rotary microtome. Sections were mounted onto 10X Genomics Xenium slides for processing, with adjacent or practice sections mounted onto SuperFrost Plus slides for histological evaluation. Slides were warmed at 44–47°C to improve adhesion and dried in a desiccator overnight before downstream processing. Adjacent sections were deparaffinized following 10X Xenium guidelines(Xenium in situ for FFPE - tissue preparation handbook) and stained with 0.05% Toluidine Blue O to evaluate section integrity, orientation, and tissue morphology prior to selecting slides for Xenium analysis. All tissue preparation and optimization leading up to and included those used for Xenium analysis were performed at the University of Minnesota.

### Xenium custom gene panel design

A custom Xenium targeted gene expression panel was designed to profile circadian and cell-type-resolved transcriptional patterns in mature Arabidopsis leaves. The panel contained 100 genes selected from core circadian clock components, clock-associated outputs, cluster biomarkers identified from the snRNA-seq time course, and literature-supported marker genes for major leaf tissues. Probe design was performed using the Arabidopsis TAIR11 genome and annotation by 10x Genomics. The complete panel is provided in Dataset S11.

### Xenium In Situ Gene Expression assay

Xenium In Situ Gene Expression was performed using the 10x Genomics Xenium platform through a Xenium RNA v1 Custom Panel, following protocols CG000580 RevE, CG000582 RevH, and CG000584 RevJ according to the manufacturer’s instructions, with the plant tissue preparation modifications described above. Briefly, FFPE sections were deparaffinized, decrosslinked, and permeabilized prior to hybridization with the custom Arabidopsis probe panel. Gene-specific probes were hybridized to target transcripts, ligated, and subjected to rolling circle amplification. Sections were counterstained with DAPI and loaded onto the Xenium Analyzer for cyclic fluorescent probe hybridization, imaging, and transcript decoding. Xenium experiments were performed by the University of Chicago at Illinois Spatial & Genome Technologies Core using a Xenium Analyzer, Xenium Software v4.0.1.4, and Xenium Onboard Analysis v4.0.1.0. Raw Xenium outputs included decoded transcript coordinates, morphology images, nuclear segmentation masks, and cell-by-gene count matrices. Detailed data processing information is outlined in the supporting information document.

### Single Nucleus Data Processing

Cell Ranger (v7.0.1) was used to generate gene by cell matrices from Demultiplexed FASTQ files by aligning with Arabidopsis TAIR10 genome. Downstream analyses were performed with Seurat (v5.1.0). Cells with low numbers of UMIs or plastid contamination were removed using Median Absolute Deviation filtering (filterbyMAD.R), and potential doublets were identified and removed through scDblFinder version 1.2(Germain et al. 2021).

After initial filtering and discarding low quality cells, the data were normalized by using the “LogNormalize” global scaling method. Following normalization, highly variable features were computed for cells of the dataset. We then used Principal Component Analysis for linear transformation and linear dimensionality reduction. After identifying the significant PCs, we implemented a graph-based clustering algorithm. Cells were embedded in a K-nearest neighbor graph structure, with cells of similar expression patterns having edges drawn to connect them followed by partitioning into communities. For clustering the cells, the Louvain algorithm was applied based on 30 top PCs.

Pseudo bulk samples were generated by aggregating the raw counts at sample and cluster level to identify oscillating gene expression. We used DeSeq2 to generate the normalized count matrix. For detecting expression of genes that oscillate throughout the time course we used METACYCLE(Wu et al. 2016). METACYCLE uses three different cycling detection algorithms i.e. meta2d: integrated ARSE, JTK-CYCLE and LS. We utilized JTK-CYCLE output to identify genes whose expression oscillates by setting cut-off Adj p-value < 0.05. The number of oscillating genes that were either specific to a cluster or shared between different clusters were illustrated by UpSet plots using the package UpSetR implemented under the *R* environment(Conway et al. 2017). JTK-CYCLE accurately measures the phase, period, and amplitude. We used phase information for each gene and calculated phase difference of genes that were shared between different clusters to identify genes that were oscillating with different phases in between different clusters.

### WGCNA analysis

To validate the phase difference observed, we used the WGCNA package(Langfelder and Horvath 2008) to analyze genes for all clusters together by including a cluster identifier appended to the gene name. A total of 13,552 genes were fed into WGCNA. To identify module based correlation, a soft threshold of 30 was used. WGCNA identified 26 modules, genes with similar expression patterns were clustered together in the same module, whereas those with different phases segregated out into different modules. We further filtered the modules based on Module Eigengene connectivity (kME), removing any gene with a kME ≥ 0.6. Additionally, to group similar modules together, we calculated the correlation matrix of the average zScores of all the modules and performed hierarchical clustering to identify major clades or expression patterns. This resulted in five modules with distinct temporal expression patterns. All code used for this analysis can be found on github (https://github.com/GreenhamLab/Arabidopsis_snRNAseq_Circadian_Transcriptome).

### scGRN Construction

Single cell GRNs were built with SCENIC implementation of GENIE3(Huynh-Thu et al. 2010; Aibar et al. 2017) by subsetting the single nucleus dataset into individual clusters and proceeding only with data from cycling genes. Several steps of the scGRN construction were adapted from the scPlant R package(Cao et al. 2023). To identify an appropriate weight threshold for each network, we shuffled and permuted our gene expression matrix into ten iterations, and identified the weight value corresponding to the 95^th^ percentile for each cluster’s aggregated network. This weight was used as a cutoff for our unshuffled networks. The same process was repeated for the Qin et al. dataset for comparative analyses. Networks were visualized and annotated in Cytoscape(Shannon et al. 2003).

### Xenium data processing and cell segmentation

Primary Xenium outputs were generated for mature Arabidopsis thaliana leaf sections collected at ZT48 and ZT60 using a custom 100-gene Xenium panel. Decoded transcripts were filtered to retain molecules with Q-score ≥ 20. Negative control probes, negative control codewords, and unassigned codewords were excluded from downstream analyses. Because mature plant leaf cells were not accurately represented by isotropic nuclear expansion alone, cell segmentation was refined using Baysor. Baysor was run on Xenium transcript coordinate files, and Baysor-derived transcript-to-cell assignments were used to generate filtered cell-feature matrices for downstream analysis.

Filtered cell-feature matrices were imported into Python using Scanpy and Squidpy. Counts were normalized to 10,000 transcripts per cell, log-transformed, and used for dimensionality reduction and clustering. Principal component analysis was performed using 20 principal components. A nearest-neighbor graph was constructed using 15 neighbors and 20 PCs, followed by UMAP embedding and Leiden clustering at resolution 0.5. Samples were analyzed independently by time point for the primary spatial clustering and spatial autocorrelation analyses.

### Cell type annotation and spatial analysis

Spatial clusters were annotated by comparing Xenium expression patterns with marker genes, cluster biomarkers, cell-type assignments from the matched single-nucleus RNA-seq circadian time course, and observation of the tissue. Epidermal, mesophyll, vascular, and guard-cell-associated spatial domains were assigned using expected marker expression patterns and snRNA-seq-derived biomarkers. Marker expression and transcript localization were manually inspected in Xenium Explorer. Spatial neighborhood relationships were calculated using Squidpy. Spatial neighbor graphs were generated from cell centroid coordinates using sq.gr.spatial_neighbors() with coord_type=“generic”. Neighborhood enrichment and co-occurrence analyses were calculated with sq.gr.nhood_enrichment() and sq.gr.co_occurrence() to identify spatial clusters enriched or depleted in local neighborhoods relative to the observed tissue organization.

Gene-level spatial autocorrelation was calculated independently for each time point using Moran’s I across segmented cells for all genes in the Xenium panel. Genes were ranked by Moran’s I within each sample, and per-sample Moran’s I tables were exported for comparison across time points. Spatial expression plots were generated from log-normalized cell-level counts and transcript coordinate overlays, with spatial axes plotted at equal aspect ratio to preserve tissue geometry.

## Acknowledgements

This work was supported by the resources and staff at the University of Minnesota Genomics Center who performed all sequencing on this project, and by the resources and staff at the University of Minnesota University Imaging Centers (UIC, SCR_020997) who assisted with single nucleus sample preparation assessment. The FFPE tissue sections were prepared at UMN and processed using the Xenium spatial transcriptomics assay by the University of Illinois Chicago (UIC) Spatial and Genome Technologies (UIC-SGT) Core, supported by the UIC Research Resources Center (UIC-RRC) and the UI Cancer Center (UIC-UICC) Shared Resources. We gratefully acknowledge Yujun Feng for her critical contributions to Xenium sample preparation, instrument operation, and high-quality data generation. We wish to thank Rui Kuang and Tianci Song for feedback on data processing approaches.

## Authors’ contributions

AZ, ZAM, and KG designed the research. AZ, ZAM, DS, AM, and KG performed the research. AZ and ZAM analyzed the data. AZ, ZAM, and KG wrote the paper.

## Availability of data and materials

All code used to process and analyze the reported data is available at https://github.com/GreenhamLab/Arabidopsis_snRNAseq_Circadian_Transcriptome. Raw sequencing data is available through SRA through BioProject accession #PRJNA1277851. Raw and processed files associated with the Xenium spatial transcriptomics, as well as the processed snRNA-seq Seurat object, have been deposited at the Gene Expression Omnibus (GEO), and will be available upon publication.

## Declaration of generative AI and AI-assisted technologies in the manuscript preparation process

During the preparation of this work, the authors leveraged Codex (5.3-mini and 5.5) and ChatGPT for code review related to data processing and to identify areas in the manuscript that could benefit from clearer language. The authors reviewed and edited the output as needed and take full responsibility for the content of the published article.

## Supplemental Tables

**Dataset S1.** Cell ranger quality control metrics for single nucleus libraries.

**Dataset S2.** Cell type specific genes.

**Dataset S3.** Gene Ontology enrichment results for snRNA-seq clusters.

**Dataset S4.** List of pseudo-bulk cycling calls.

**Dataset S5.** Cluster-resolved cycling gene calls.

**Dataset S6.** WGCNA temporal clade assignments for cluster-resolved rhythmic genes.

**Dataset S7.** Cluster-resolved single-cell gene regulatory network edges.

**Dataset S8.** Qin et al. single-nucleus gene regulatory network edges used for comparison.

**Dataset S9.** CCA1, LNK2, and PRR7 target specificity across broad cell types.

**Dataset S10.** Gene Ontology enrichment for CCA1/LNK2/PRR7 target specificity groups.

**Dataset S11.** Xenium 100-gene panel and transcript detection across spatial samples.

**Dataset S12.** Xenium sample, segmentation, and cell-cluster summary metrics.

**Dataset S13.** Xenium cluster marker genes.

**Dataset S14.** Cell type-resolved Xenium expression values for core clock genes.

**Dataset S15.** Moran’s I spatial autocorrelation results for Xenium genes.

