## Supplementary figures and images for "Tissue Context Shapes the Circadian Transcriptome of the Arabidopsis Leaf"

### Figure S1

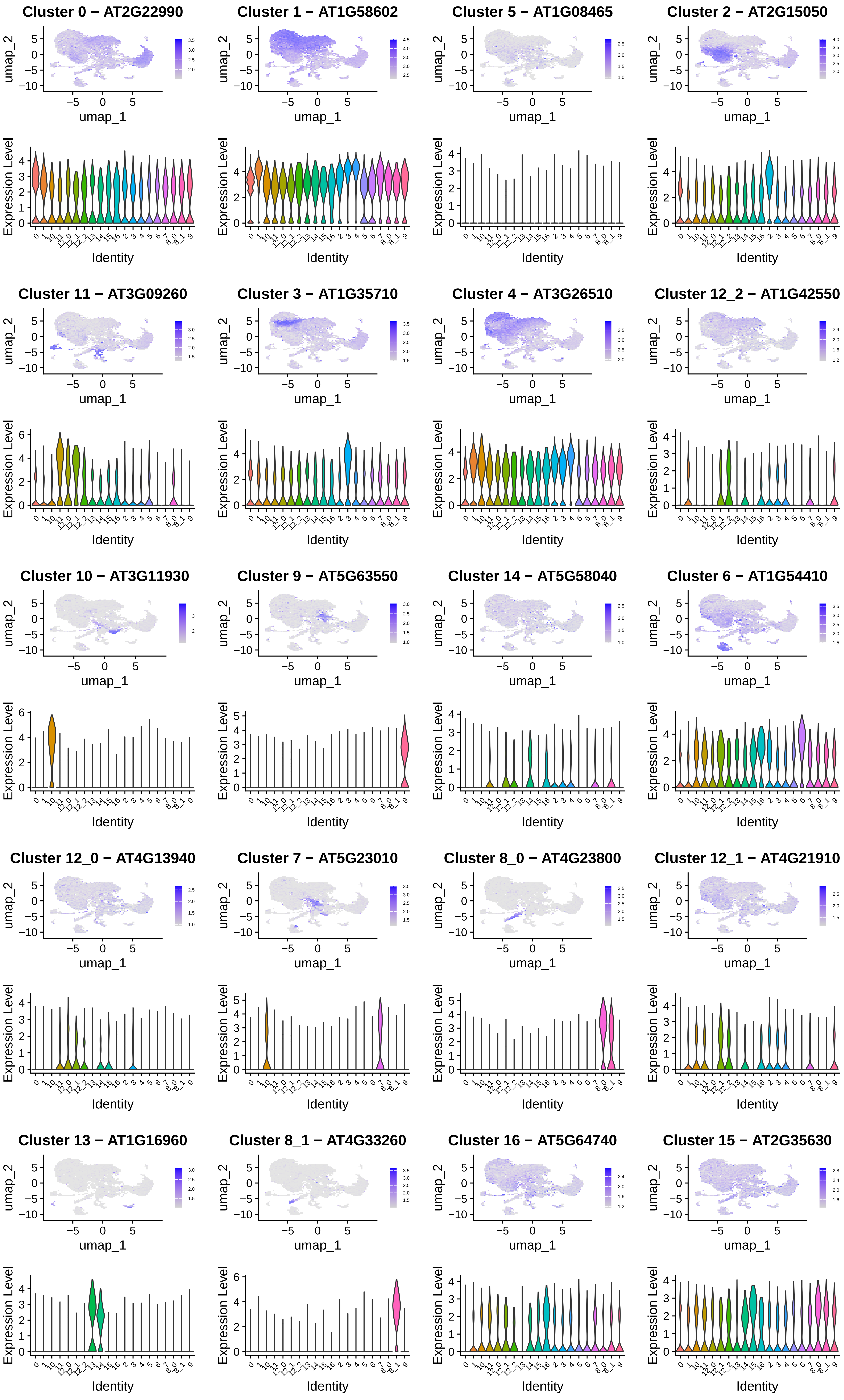

### Figure S2

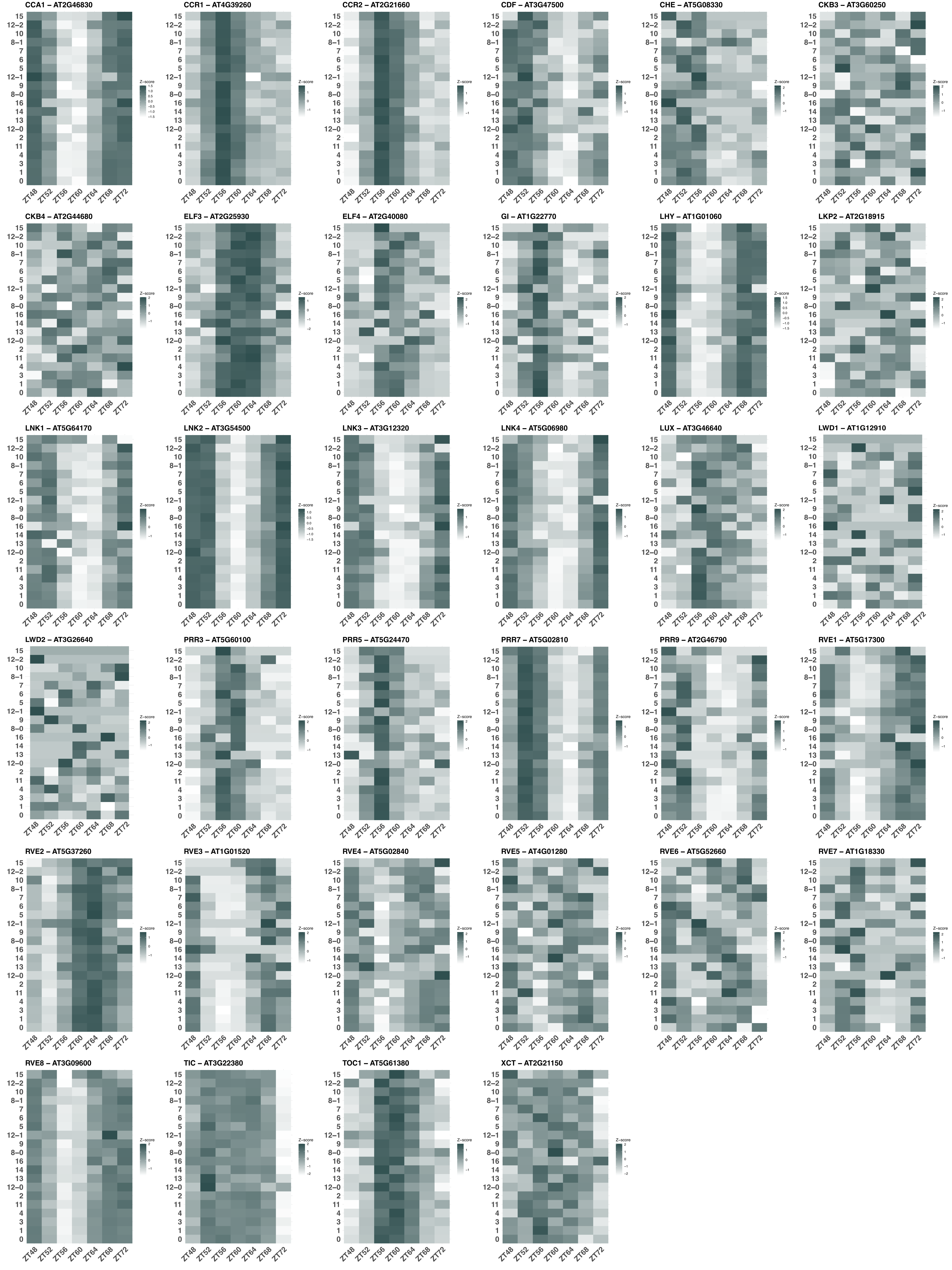

### Figure S3

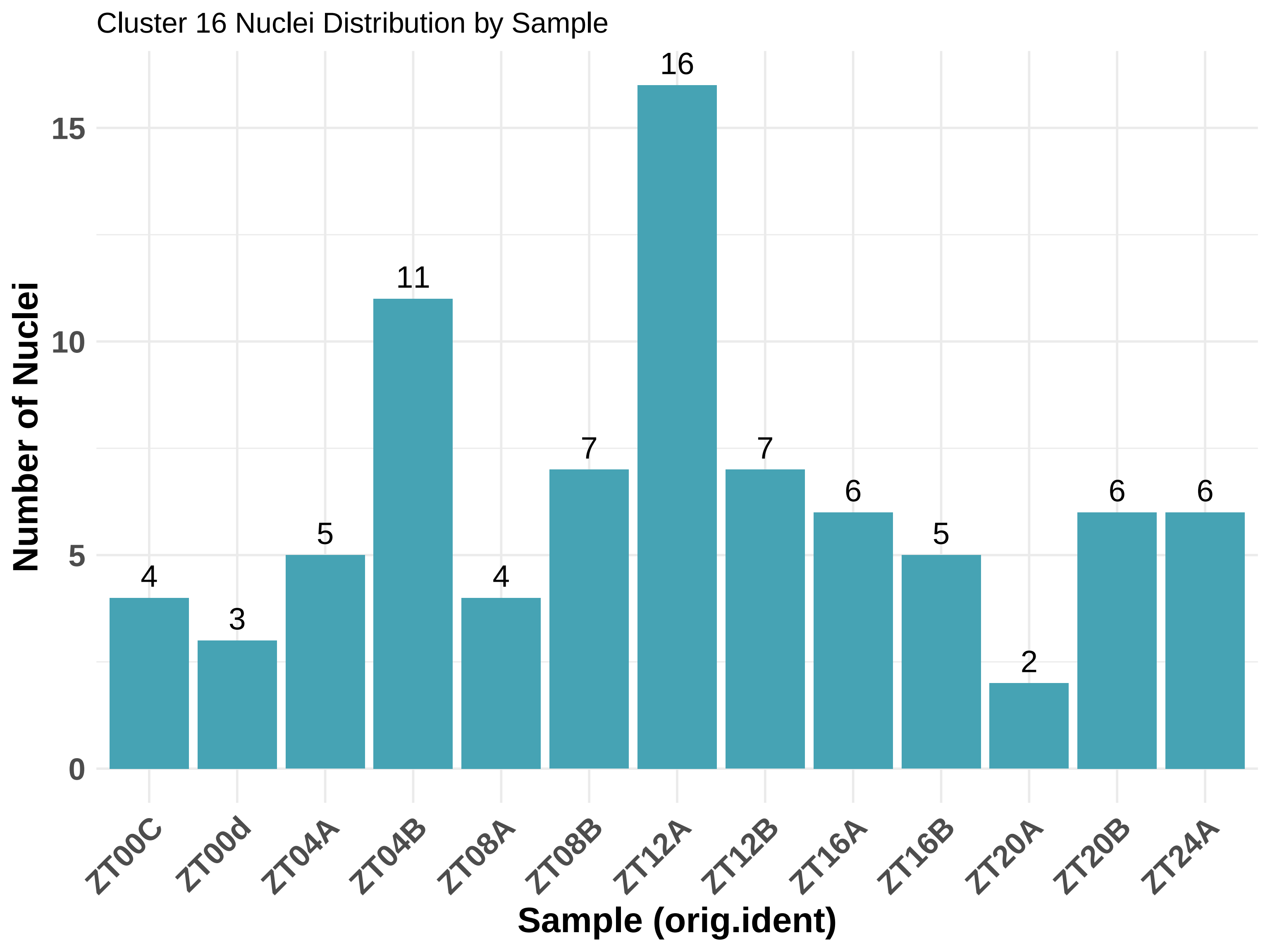

### Figure S4

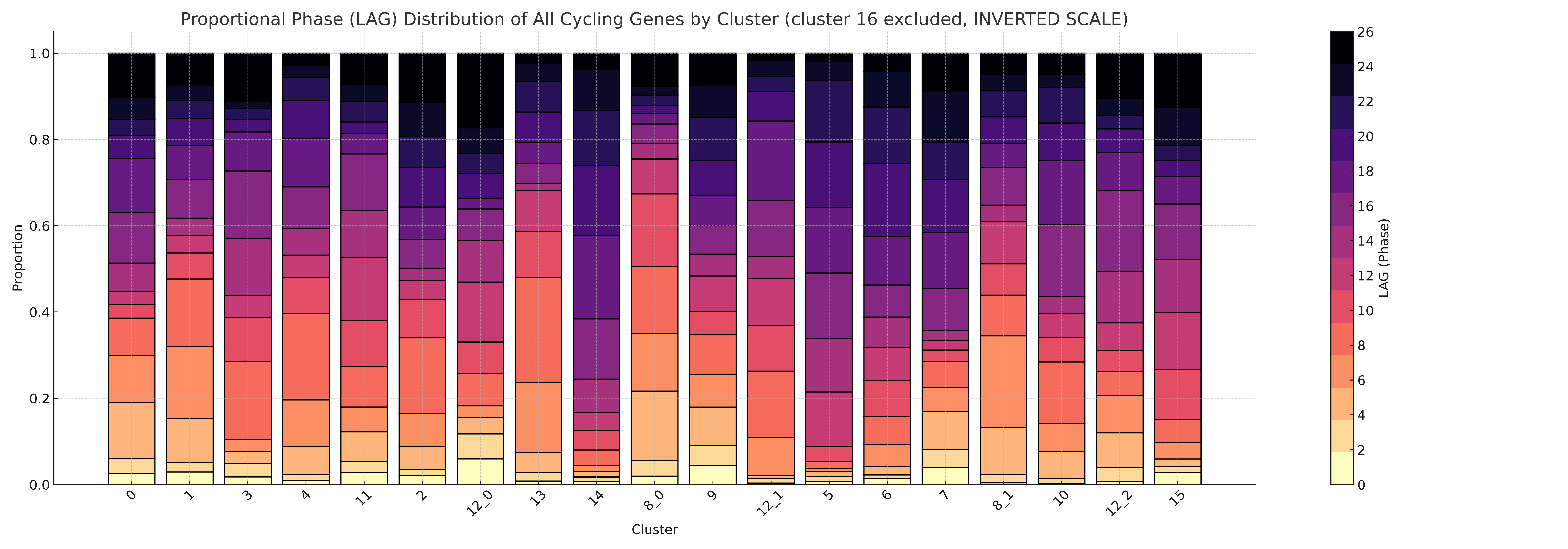

### Figure S5

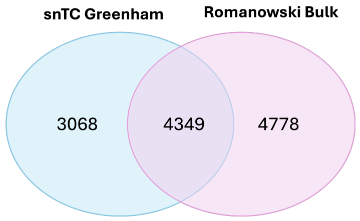

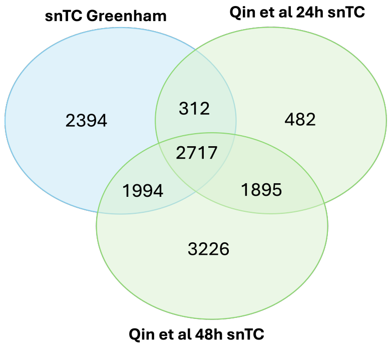

### Figure S6

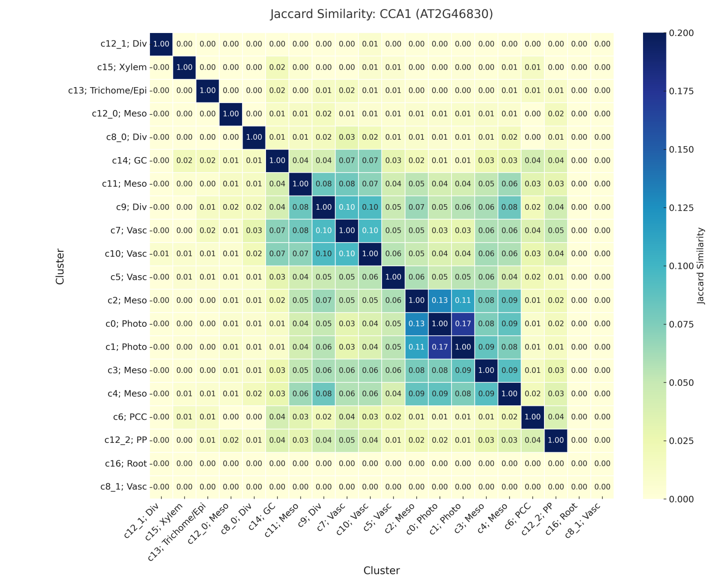

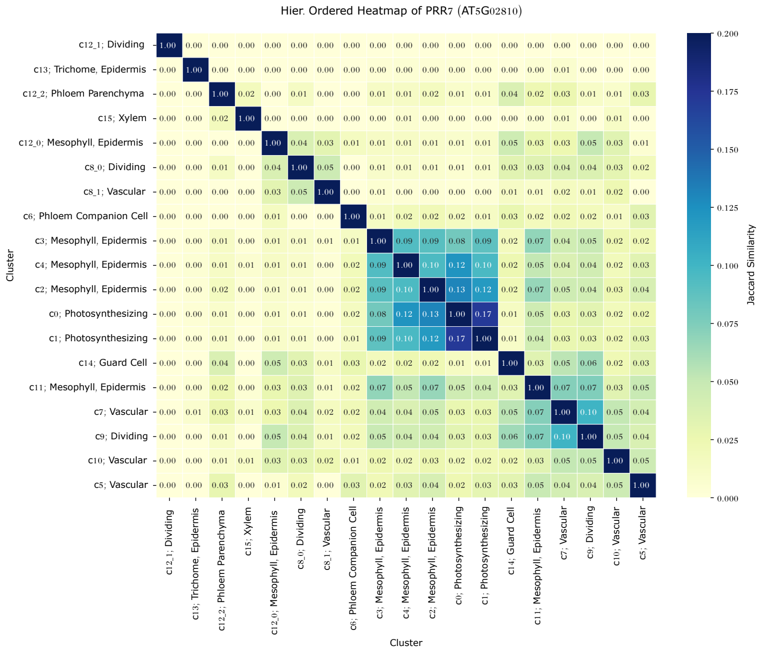

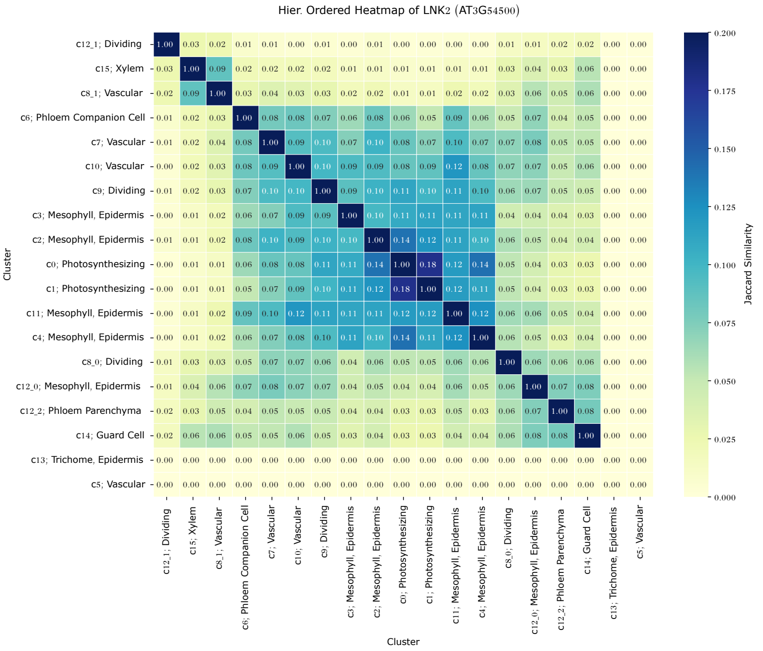

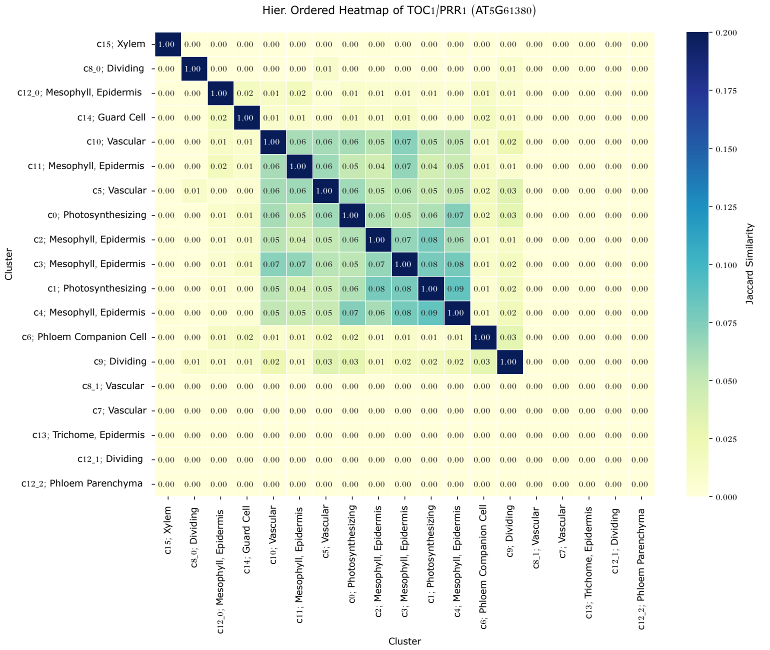

### Figure S7

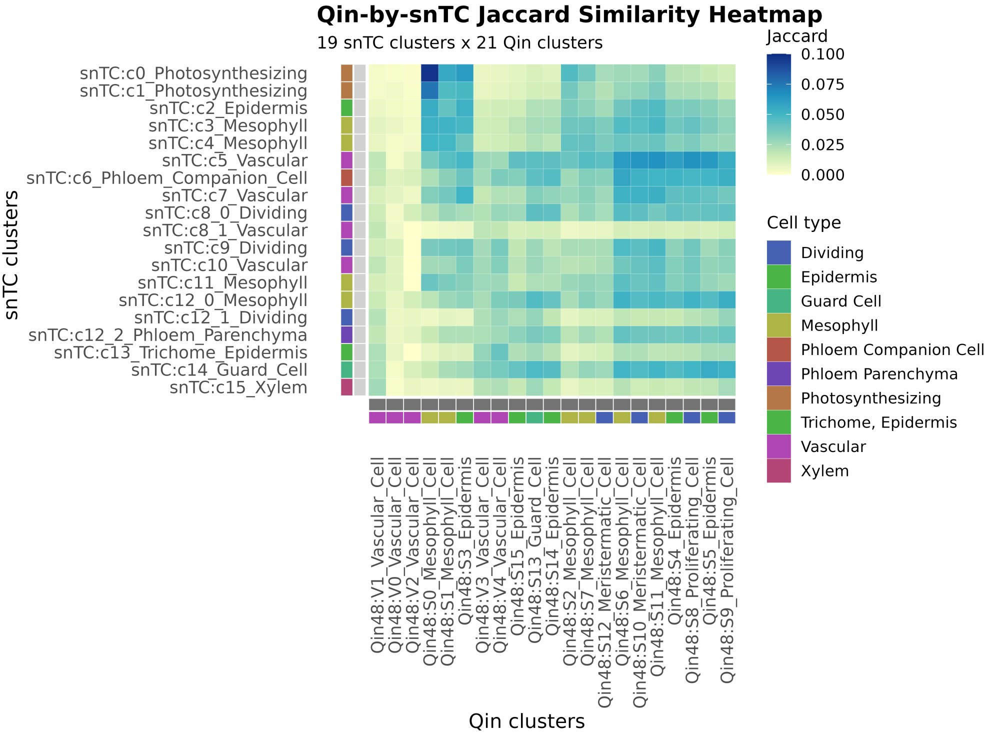

### Figure S8

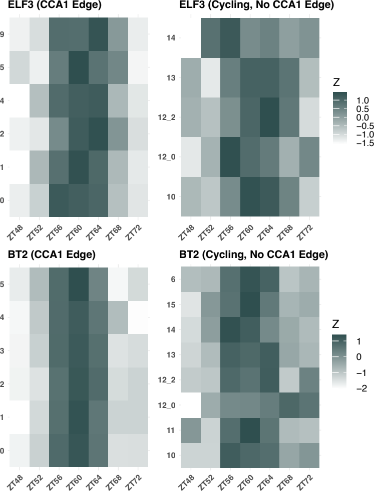

### Figure S9

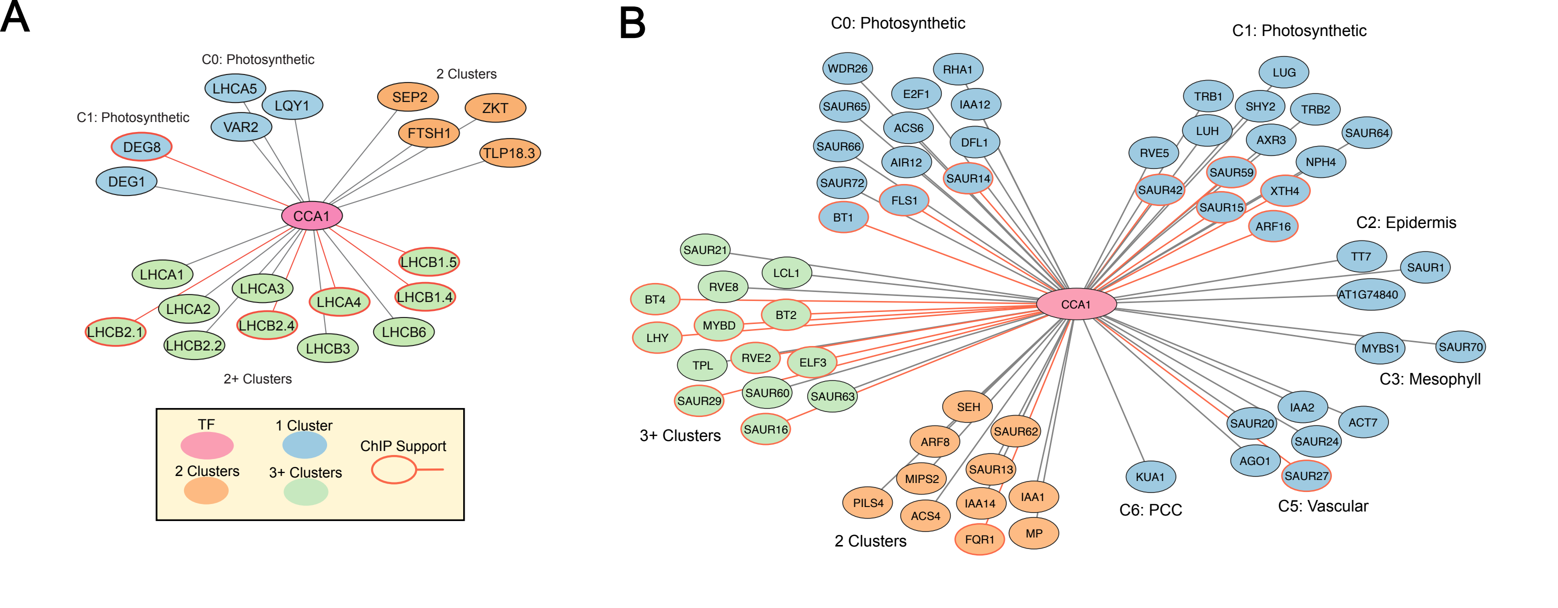

### Figure S10

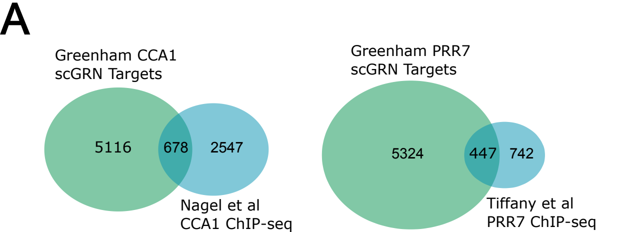

### Figure S11

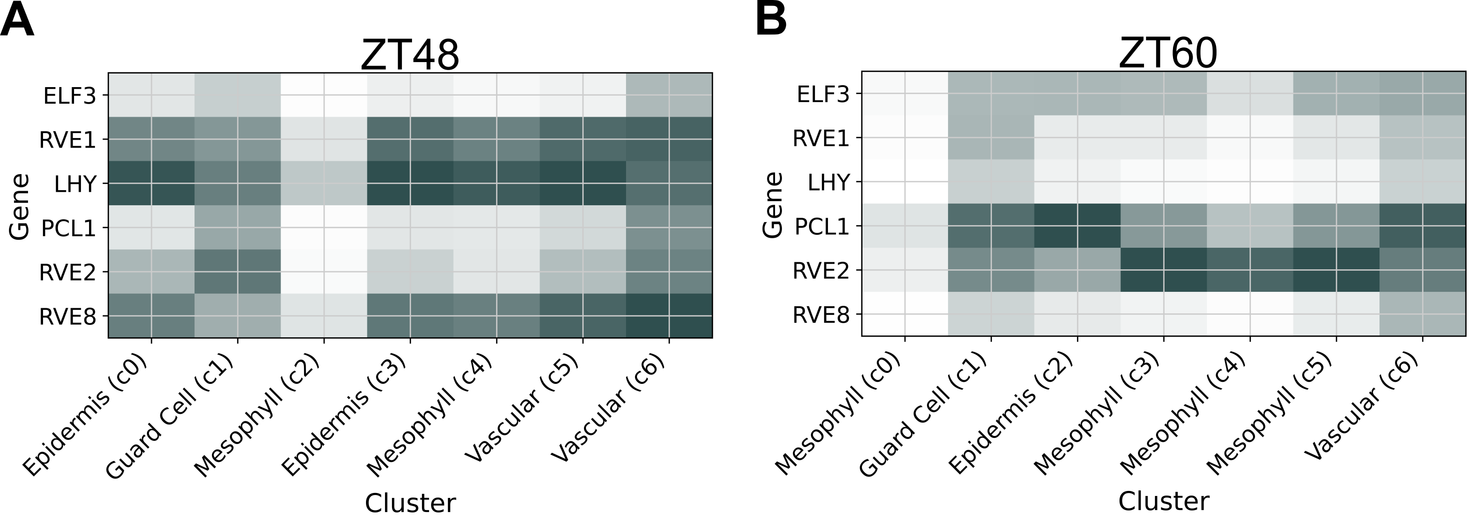
